# Model-based EEG phenotyping uncovers distinct neurocomputational mechanisms underlying learning impairments across psychopathologies

**DOI:** 10.1101/2025.11.01.685815

**Authors:** Nadja R. Ging-Jehli, Rachel Rac-Lubashevsky, Krishn Bera, Alyssa Roberts, Allie Loder, Megan A. Boudewyn, Cameron S. Carter, Molly A. Erickson, James M. Gold, Steven J. Luck, J. Daniel Ragland, Andrew P. Yonelinas, Angus W. MacDonald, Deanna M. Barch, Michael J. Frank

**Affiliations:** Carney Institute for Brain Science, Department of Cognitive and Psychological Sciences, Brown University, Providence, RI; Department of Psychology, University of Minnesota, MN; Department of Psychology, University of California, Santa Cruz, CA; Department of Psychology, University of California, Irvine, CA; Department of Psychiatry & Behavioral Neuroscience, University of Chicago; Department of Psychiatry, University of Maryland School of Medicine, Baltimore, MD; Department of Psychology, University of California, Davis, CA; Department of Psychology, Washington University in St. Louis, MO

**Keywords:** computational phenotyping, computational psychiatry, reinforcement learning, working memory, decision-making, transdiagnostic, depression, bipolar, schizophrenia

## Abstract

**Background:** Major depressive disorder (MDD), bipolar disorder (BP), and schizophrenia (SCZ) involve learning impairments with poorly understood mechanisms. Understanding both the similarities and differences in these mechanisms is important to guide the development of new, targeted interventions.

**Methods:** 255 participants diagnosed with MDD (n=54), BP (n=47), SCZ (n=67) or without any diagnoses (CTRL; n=87) performed an associative learning task. Computational modeling quantified the mechanistic interplay between working memory (WM) and reinforcement learning (RL). The latent RL and WM signatures in the EEG dynamics showed shared and distinct neurocognitive mechanisms underlying learning.

**Results:** All clinical groups showed learning impairments at the behavioral level. Model-based EEG analyses linked these impairments to distinct patterns in the dynamic interplay between latent RL and WM mechanisms, contrasting with the typical patterns observed in CTRL. SCZ was characterized by reduced neural markers of WM, weakening the cooperative influence of WM onto RL (reduced WM recruitment), and reduced integration of negative feedback. Conversely, MDD was characterized by reduced reciprocal influence of RL onto WM, reducing the tendency to upregulate WM contribution with reward history (impaired WM management). Finally, BP was characterized by deficits in both WM and RL recruitment, along with higher WM decay.

**Conclusions:** Behavioral learning impairments that appear similar across clinical groups can be linked to distinct neurocognitive mechanisms via integrative neurocomputational modeling. Our approach provides insights into the interplay of underlying learning mechanisms and how they manifest differently across psychopathologies.

**Citation:** This manuscript is a preprint version of the later manuscript accepted for publication in **Biological Psychiatry: Global Open Science**. The content may differ from the final published version following peer review and editorial revisions.

Ging-Jehli, N.R., Rac-Lubashevsky, R., Bera, K., Roberts, A., Loder, A., Boudewyn, M.A., Carter, C.S., Erickson, M., Gold, J., Luck, S.J., Ragland, J.D., Yonelinas, A.P., MacDonald III, A.W., Barch, D.M., & Frank, M.J. (2025). Model-based EEG phenotyping uncovers distinct neurocomputational mechanisms underlying learning impairments across psychopathologies. Preprint at bioRxiv.

## Introduction

Learning impairments are prevalent across mental health conditions, yet their underlying mechanisms remain inconclusive.^1,2^ Cognitive neuroscience and computational psychiatry offer approaches to dissect latent mechanisms underlying learning and decision-making processes more broadly.^3–10^ Ultimately, examining how neurocognitive, motivational, and situational mechanisms interact at different timescales could build an understanding of how their interplay reinforces distinct symptom patterns across psychopathologies.^3–5,11^ This lays the foundation for identifying and predicting when and which mechanisms should be targeted in interventions which is useful particularly for conditions with complex treatment needs like major depressive disorder (MDD), bipolar disorder (BP), and schizophrenia (SCZ).

Towards building a comprehensive mechanistic understanding of symptom profiles, it is valuable to first evaluate whether mechanistic models can effectively differentiate transdiagnostic features that may appear identical at the behavioral level. A promising approach is to focus on a specific context known for its sensitivity to clinical challenges, such as learning impairments. This paves the way for more comprehensive investigations in the future (e.g., symptom manifestations across contexts). Past research emphasizes the importance of complementing computational models with experimental paradigms that are sensitive to differentiate between model-simulated mechanisms; and grounding them both in neuroscientific theory.^3,9^ Here, we demonstrate how such a mechanistic approach separates latent neurocognitive mechanisms of learning impairments; and by doing so identifies shared and distinct characteristics across MDD, BP, and SCZ.

The goal of this study is to characterize the nature of cognitive impairments across disorders using a mechanistic framework that dissociates reinforcement learning and working memory contributions to behavior. For doing so, we focus on instrumental learning, which refers to the formation of stimulus-response associations via operant conditioning, and which relies on multiple neurocomputational mechanisms modulated by dopaminergic dynamics in distinct prefrontal-basal ganglia pathways.^12–14^ These mechanisms can be effectively studied through neurocognitive frameworks of reinforcement learning (RL) and working memory (WM).^8,15^ Individuals with MDD, BP, and SCZ often demonstrate impairments in instrumental learning.^6–8^ Multiple neurocognitive mechanisms contribute to learning and the frameworks of reinforcement learning (RL) and working memory (WM) can disentangle these processes. ^12,16^ Combining RL and WM frameworks into the same model acknowledges that effective associative learning requires both integrating the consequences of past actions (RL) and maintaining information (i.e., stimulus-action-outcome associations) over short periods (WM). Accounting for interactions between RL and WM processes within the same model differentiates the underlying sources of deficits when combined with tasks that systematically target these mechanisms across different task conditions.^4,17^ The RLWM task has been shown to satisfy these requirements, producing behavioral learning curves consistent with the predictions of the mechanistic model in multiple (mostly non-clinical) studies.^12,15,18–20^ Prior work has shown that this approach can refine the clinical understanding of underlying cognitive deficits.^10,14^

In this study, we identified the unique neurocognitive learning dynamics between individuals with MDD, BP, SCZ, and those without clinical diagnoses (CTRL) for the first time (Table 1). Importantly, combining the mechanistic RLWM task^12^ with established model-based EEG analyses^15,18^ allowed for: 1. validating the latent neurocognitive dynamics predicted by the computational model; 2. tracing individual differences in real-time learning dynamics; and 3. assigning cognitive interpretation to the extracted physiological measures. Although clinical groups exhibited similar behavioral impairments, we found that the model-based EEG analyses linked these impairments to alterations in different neurocomputational components. We next summarize key characteristics of the task and model-based analyses.

**Table 1.**
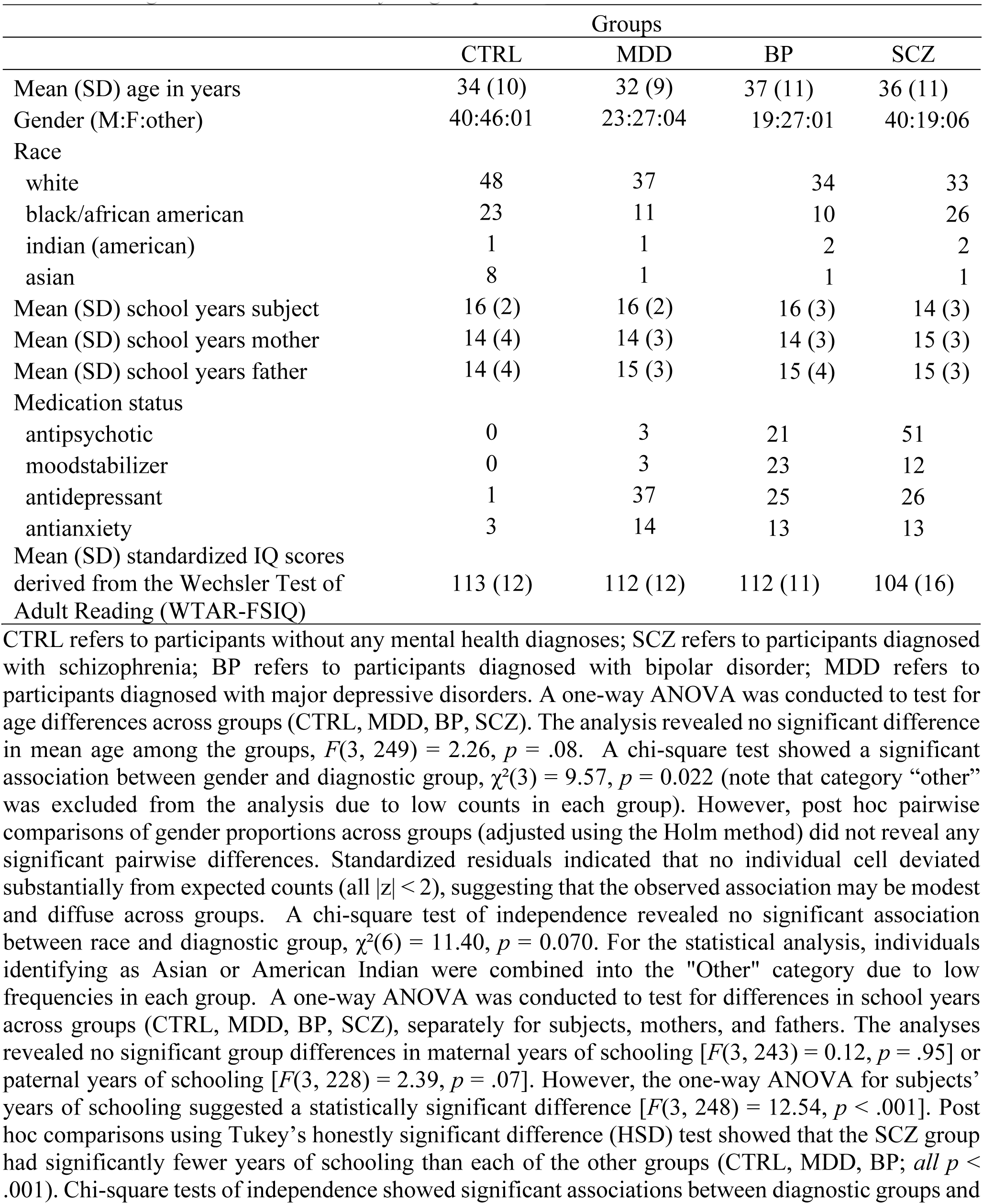
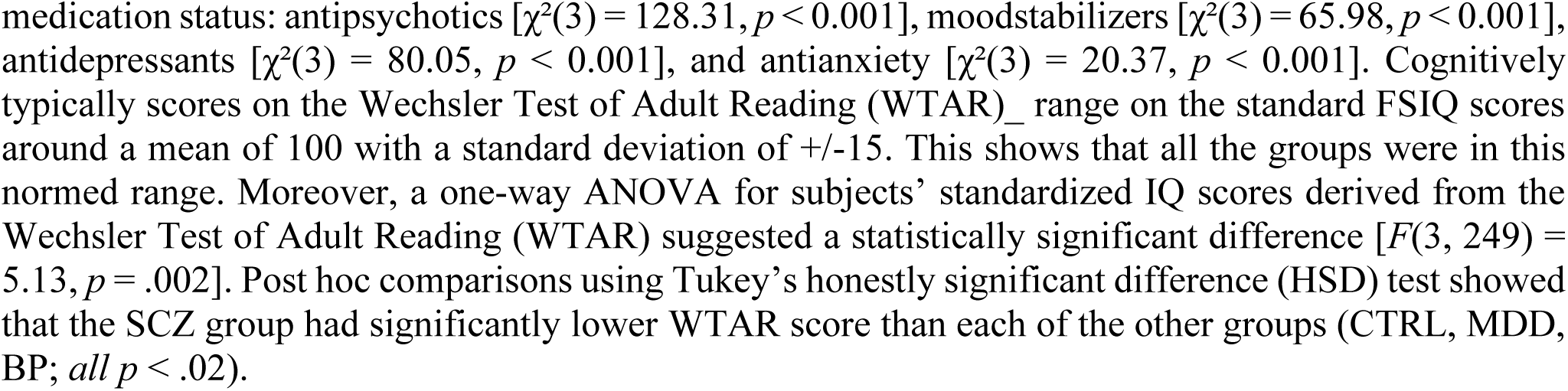
Background characteristics of the groups.

|  | Groups |  |  |  |
| --- | --- | --- | --- | --- |
|  | CTRL | MDD | BP | SCZ |
| Mean (SD) age in years | 34 (10) | 32 (9) | 37 (11) | 36 (11) |
| Gender (M:F:other) | 40:46:01 | 23:27:04 | 19:27:01 | 40:19:06 |
| Race |  |  |  |  |
| white | 48 | 37 | 34 | 33 |
| black/african american | 23 | 11 | 10 | 26 |
| indian (american) | 1 | 1 | 2 | 2 |
| asian | 8 | 1 | 1 | 1 |
| Mean (SD) school years subject | 16 (2) | 16 (2) | 16 (3) | 14 (3) |
| Mean (SD) school years mother | 14 (4) | 14 (3) | 14 (3) | 15 (3) |
| Mean (SD) school years father | 14 (4) | 15 (3) | 15 (4) | 15 (3) |
| Medication status |  |  |  |  |
| antipsychotic | 0 | 3 | 21 | 51 |
| moodstabilizer | 0 | 3 | 23 | 12 |
| antidepressant | 1 | 37 | 25 | 26 |
| antianxiety | 3 | 14 | 13 | 13 |
| Mean (SD) standardized IQ scores<br>derived from the Wechsler Test of<br>Adult Reading (WTAR-FSIQ) | 113 (12) | 112 (12) | 112 (11) | 104 (16) |

In the RLWM task, participants learn stimulus-response associations through trial-and-error and subsequent feedback (Fig. 1A). The task is structured into multiple blocks of trials, parametrically manipulating WM demand. Specifically, WM demand varies between blocks due to differences in *set size* (the number of presented stimulus-response associations) and within blocks due to *delay* (the number of intervening trials before the same stimulus-response association is repeated). RL contributions are indexed by so-called learning curves of presented stimulus-response associations. Specifically, the increase in accuracy over time as a function of correctly repeated responses to a given stimulus-response association (*reward history*). Further details on task structure and specific manipulations are in the Methods section.

**Figure 1.**
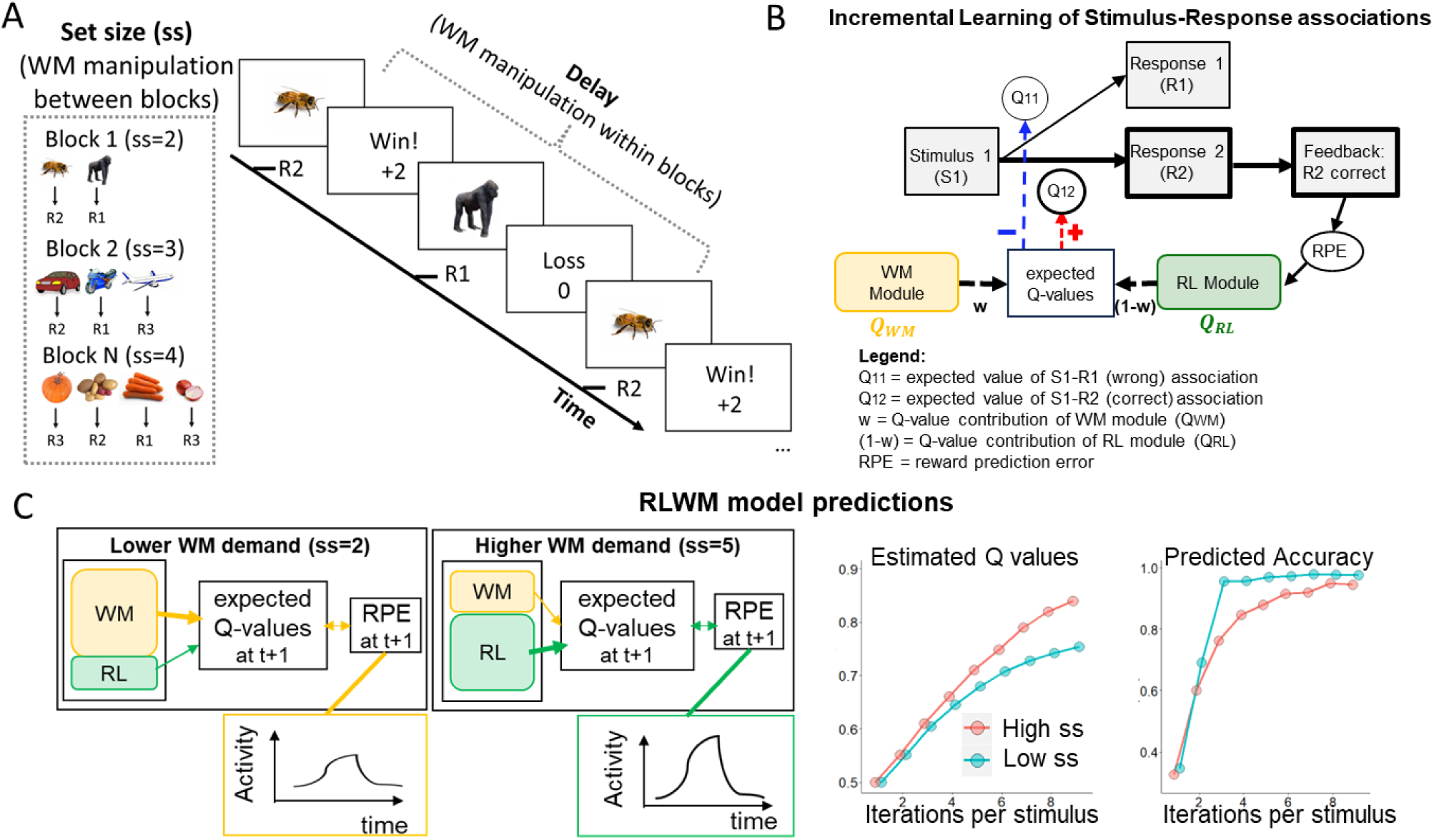
Characteristics of the RLWM task and its computational model. **A.** Participants learn associations between stimuli and three response options via feedback. Shown is the procedure of one task trial. WM load is parametrically manipulated with between-block manipulation of set size (ss) and within-block manipulation of delay. **B.** The mechanistic computational model simulates the dynamic interaction of latent WM and RL components that contribute to the Q-value-based learning process across trials. RPEs used for updating RL Q values are informed by RL expectations Q_RL_ but also cooperative impact of WM expectations (Q_WM_), weighted by the individual’s reliance on WM. Selected model parameters are summarized in the pink box. **C.** Selected key model predictions, emulating how both WM and RL processes contribute to expected Q values and reward prediction errors (RPEs). Under lower WM demand, expected Q values can be retained in working memory, which diminishes RPEs in early learning stages. Conversely, under higher WM demand, working memory is above capacity, increasing the relative contribution of RL-based computations and leading to higher RPEs and RL Q values, despite worse accuracy. The predicted cooperative effect of RL and WM processes on estimated Q values and accuracy are displayed in the bottom subplots, respectively. These subplots are recreated from past studies.^15,18^

The RLWM model is fitted to trial-by-trial choice data to capture the observed learning curves. The model includes a faster (but capacity-limited) WM module as well as a slower (but capacity-free) RL module. Each module contributes to the formation of an expected value (*Q-value*) for each stimulus-response association. After presenting a stimulus, the response with the highest Q-value is selected (Fig. 1B; Table 2). The WM module facilitates rapid information processing thought to be mediated by active maintenance of stimulus-action-outcome contingencies within prefrontal cortex.^21–24^ Capacity constraints make the WM module vulnerable to load and interference effects. Conversely, the slower RL module, thought to rely on corticostriatal synaptic plasticity, gradually strengthens learned associations through trial-by-trial feedback-driven reward prediction error (*RPE*) via the classical Q-learning algorithm.). As such the expected reward value (*Q-value*) of the chosen stimulus-response association increases progressively with accumulated reward history, gradually easing the computational load initially handled by the faster, capacity-limited WM module.

**Table 2.**
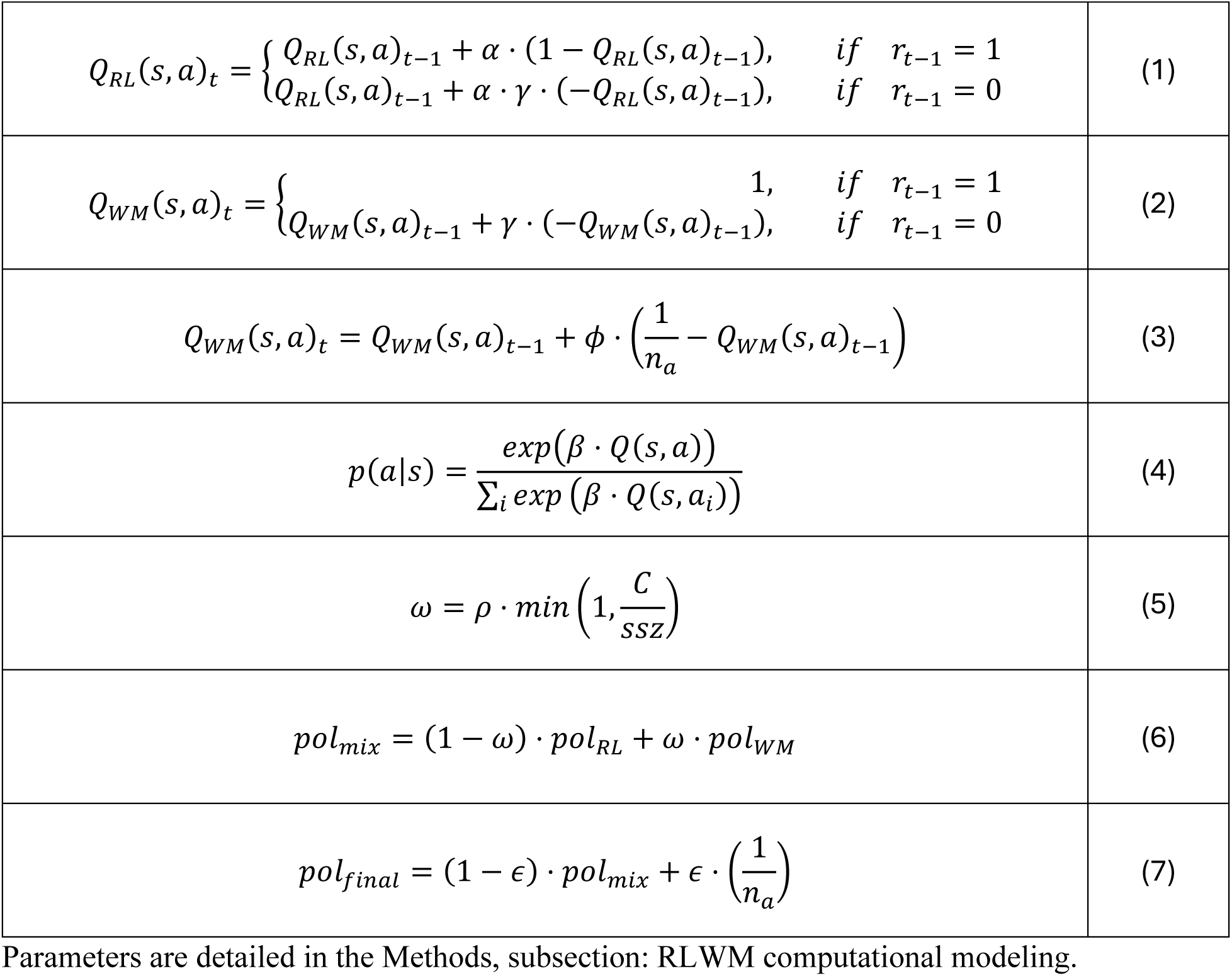
Equations specifying the RLWM computational modeling approach.

Initial work with this model assumed that the two modules were independent, in part because individual variations in RL and WM parameters were associated with genetic components affecting striatal plasticity and prefrontal function, respectively.^12^ More recent studies have shown, however, WM and RL systems also interact *cooperatively*.^14,15,18,19^ In particular, these studies have shown that the RPEs needed for learning by the RL system are influenced by expectations that are held in WM. Consider, for example, a trial in which participants respond correctly and get positive feedback. When stimulus-outcome associations can be held in WM (during low WM load), this correct outcome will match WM-based expectations, reducing the RPE experienced by the RL module (Fig. 1C). Conversely, under high WM demand, outcomes are not reliably expected in WM, and accordingly, larger RPEs cause increased Q-value updating in RL, reducing the burden on the capacity-limited WM module. This cooperative tradeoff is supported by fMRI, EEG and behavioral studies.^14,15,18,19^ Importantly, these predictions do not imply that *either* RL *or* WM processes are recruited on a given trial but rather they interact dynamically within and across trials. This phenomenon provides an opportunity to study how these interactions might vary in the clinical conditions studied here.

To extract neural signatures of WM and RL components, and how they interact, we employed an established model-based EEG approach and focused on correct responses (detailed in the Methods).^15,18^ To summarize key aspects of this approach, fitting the RLWM model to behavioral data provided trial-by-trial measures of latent RL-based computations including Q-values and RPEs. These model variables, together with WM manipulations (set size and delay), subsequently served as blueprints to extract trial-wise neural RL and WM markers. The obtained markers consist of spatiotemporal EEG patterns quantifying the degree to which participants engage in WM and RL computations at any given time and brain (surface) location.^15,18^ For example, the neural RL marker reflects brain correlates of the stimulus-response expectation within the RL system, providing a neural learning curve across stimulus iterations. Conversely, the neural RPE marker reflects individuals’ surprise following successful outcomes, which declines over trials as associations are learned. Neural WM markers reflect the degree to which a given stimulus elicits brain activity related to the need to manage multiple concurrent stimuli in WM (see Methods for more details). We will demonstrate that, although behavioral analyses indicate similar learning impairments across clinical groups, the sources of these changes can be differentiated through this integrative model-based EEG analyses.

To preview the main findings, all three clinical groups showed poorer performance as compared to CTRL. First, BP and MDD were both characterized by poorer learning under higher than lower WM demand (i.e., higher set size effects). The model-based EEG analyses attributed these observable learning impairments to distinct underlying mechanisms. While BP was characterized by higher WM decay, MDD was characterized by over-reliance on WM especially when it would have been beneficial to rely on RL (i.e., reduced effect of reward history on WM recruitment). We refer to this pattern as a deficit in WM “management” because it reflects the disruption of an adaptive process putatively related to frontostriatal-WM gating.^25^ Conversely, SCZ was characterized by WM impairments already under lower demand and reduced influence of negative compared to positive feedback on learning, replicating earlier effects in this task^10^ (but see below for novel EEG findings indicating alterations in cooperative WM-RL interactions). In summary, all three clinical groups showed different patterns of cooperative WM-RL interactions compared to CTRL, which were only detectable through the model-based EEG analyses. We will subsequently unpack these main findings.

## Results

255 participants without mental health diagnoses (CTRL, n=87) or with schizophrenia (SCZ, n=67), major depressive disorder (MDD, n=54) or bipolar disorder (BP, n=47) performed the RLWM task during EEG recording (Table 1). In the first part of this section, we summarize findings from analyses including all participants, highlighting the dissociable computations related to WM and RL in both behavioral and EEG measures and replicating past results.^15,18,19^ The general findings build the foundation for interpreting clinical similarities and differences in the second part. To dissociate WM and RL effects on performance measures, we used mixed-effect regressions whose complete outputs are provided in the Supplement.

### Part I: General Findings Across All Participants

Participants incrementally learned the stimulus-response associations, as demonstrated by the learning curves (Fig. 2A) that increased with reward history [*β* = 0.78, *SE* = 0.04, *p* < 0.001]; a proxy for latent RL-based computations.^10,14,15,18,19^ Fig. 2A also shows that the RLWM model (introduced earlier) captured these learning curves well.

**Figure 2.**
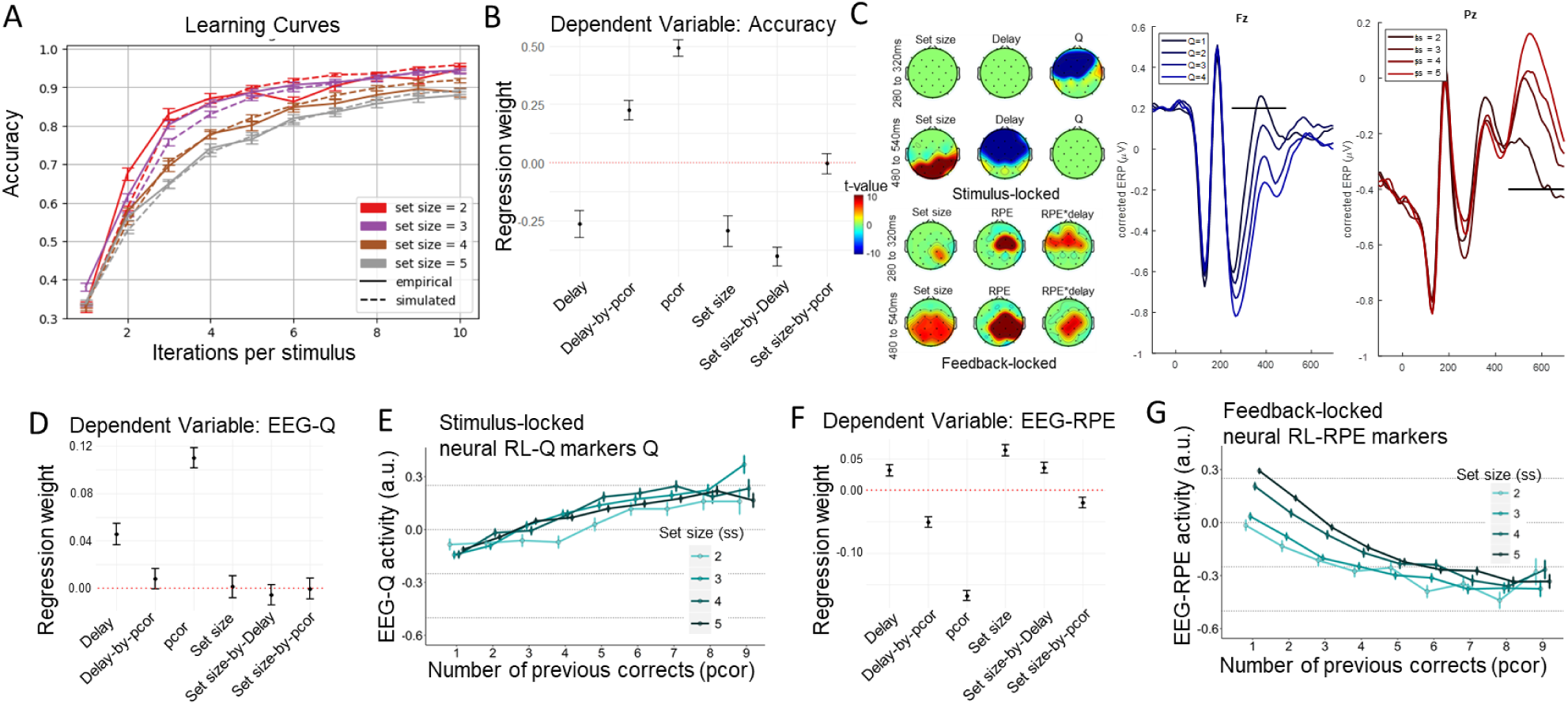
Analyses across all participants replicate previous RLWM behavioral, EEG and mechanistic modeling findings. **A.** Accuracy (averaged across trials and participants) curves by number of stimulus iterations show continuous learning over time that varies with set size. Vertical bars represent within-subject SEs. **B.** Coefficient weights from logistic mixed-effect regressions with trial-based accuracy as dependent variable and covariates: set size, delay, reward history (pcor) and their interactions. This shows dissociable contributions of WM components (delay, set size, and their interactions) and incremental RL (effect of previous corrects). See Suppl. Table S1 for regression output. **C.** EEG-based markers of WM and RL during decision-making and feedback-processing. The topographic scalp maps illustrate the effects of each predictor (set size, delay, Q) across an early (top row) and later (bottom row) time window. The maps are presented for stimulus-locked events (left plots) and for feedback-locked events (right plots). These spatiotemporal patterns induced by GLM can also be visualized by classical ERPs correcting for other terms in the GLM. Here, corrected ERP plots show the effect of the two main predictors discussed in the Result section (see Suppl. Fig. S1 for analogous plots of all predictors), namely: RL markers in blue (by RL value quartiles) and neural set size markers in red (by set size) on the voltage of significant electrodes (Fz and Pz both at central lines). Horizontal black lines reflect significant time points after permutation correction. **D.** Coefficient weights from mixed-effect regressions with neural RL markers as dependent variable and predictors: set size, delay, reward history (pcor), and their interactions. As expected, pcor and delay contributed to neural RL markers consistent with past studies, showing more RL recruitment under longer delay.^15,18^ See Suppl. Table S2 for regression output. **E.** Neural RL markers, indexing incremental RL, increased with reward history (pcor). Vertical bars represent within-subject SEs. Indices were extracted from trial-by-trial model-based EEG analyses using trial-based Q-value estimates as a blueprint. **F.** Coefficient weights from regressions with neural RPE markers as dependent variable and predictors: set size, delay, reward history (pcor), and their interactions. As expected, neural RPE was inversely related to pcor but positively related to set size. See Suppl. Table S3 for regression output. **G.** Neural RPE markers decreased with reward history (pcor). Vertical bars represent within-subject SEs. Indices were extracted from model-based EEG analyses using trial-based RPE estimates as a blueprint.

### Instrumental learning mitigates the negative performance effects of higher WM demand

We manipulated WM demand by varying set size (number of stimulus-response associations) across blocks, and delay (number of trials between successive presentations of the same stimulus) within blocks. As expected, trial-by-trial accuracy significantly decreased under higher WM demand (Fig. 2B). However, these detrimental effects diminished with accumulated reward history, demonstrating the expected positive contribution of RL to task performance.^10,14,15,18,19^

### Model-based neural markers dissect WM and RL contributions

As described earlier (see also Methods), the model-based EEG analyses used the computational parameters to extract neural markers of RL and WM components. By identifying clusters of neural activity associated with these components (see Methods), we tracked the dynamic evolution of these latent RL and WM components at a neural level over time and across trials. Fig. 2C illustrates the spatial distribution of these neurocomputational markers within the temporal windows where their association was significant. The neural RL marker of Q-values (subsequently referred to as *neural RL markers*) produced early frontal activity during decision-making (∼300ms post-stimulus), while the neural WM markers showed later parietal (for set size) and frontal (for delay) activities (∼500ms post-stimulus). Hence, RL components contributed to earlier learning, whereas WM components contributed to later stages.^10^

### Determinants of neural RL markers

As expected from RL frameworks, greater neural RL markers were related to higher accuracy during the asymptotic phase, as indicated by a significant interaction between RL marker and reward history [*β* = 0.045, *SE* = 0.014, *p* = 0.002]. Fig. 2D provides the corresponding regression weights (see figure note for model specifications). As successful experiences accumulated (reward history), participants transitioned from WM to RL computations, consistent with RLWM model predictions (Fig. 2E).^10,12,14,15,18,19^

To separate WM and RL contributions, we estimated a linear mixed-effects regression with neural RL markers as dependent variable and predictors including behavioral and neural WM components (set size, neural set size markers), RL components (reward history), and their interactions (Suppl. Table S4) This revealed positive main effects of both reward history [*β* = 0.012, *SE* = 0.007, *p* < 0.001] and set size [*β* = 0.052, *SE* = 0.006, *p* < 0.001], implying increased RL engagement not only under asymptotic learning but also under higher WM demand. We also found a negative main effect of neural set size marker [*β* = -0.33, *SE* = 0.007, *p* < 0.001], suggesting that neural markers of WM are indicative of reduced RL engagement over and above the manipulated WM demand (i.e., objective set size). Lastly, we found a significant interaction between reward history and neural set size marker [*β* = 0.011, *SE* = 0.005, *p* = 0.021]. As reward history accumulated, greater neural set size markers were related to enhanced RL computations, which are especially beneficial under higher WM demand (i.e., higher set sizes) as detailed next.^15,18^ We will later demonstrate that MDD specifically is associated with impairments in this interactive RL-WM mechanism.

### Determinants of neural set size markers

Theoretical models of frontostriatal circuits suggest that not only does WM influence instrumental learning, but RL can also reciprocally influence gating of information into and out of WM.^25^ Indeed, recent studies have shown that RL processes support adaptive *management* of WM content with experience, increasing effective WM capacity.^26^ Accordingly, we found that higher neural set size markers not only reflected WM *challenges* (due to its positive associations with WM demand as indexed by set size) but also *efficiency* in WM management (due to its positive associations with RL processes as indexed by reward history). This is evidenced by a linear mixed-effects regression (Suppl. Table S5), showing that neural set size markers increased with both set size [*β* = 0.117, *SE* = 0.004, *p* < 0.001] and reward history [*β* = 0.023, *SE* = 0.004, *p* < 0.001], along with a significant interaction between them [*β* = 0.011, *SE* = 0.004, *p* = 0.015]. As participants accumulate reward history, they form stronger stimulus-response associations within RL, freeing the WM system to efficiently prioritize how to manage its resources under larger set sizes. This highlights the need for efficient WM functioning even when the system transitions from WM to RL with learning.

We next examined whether neural set size markers predicted accuracy, beyond the effects of manipulated set size and reward history (Suppl. Table S6). Indeed, higher neural set size marker predicted higher accuracy under higher WM demand as indicated by a positive interaction between neural set size marker and set size [*β* = 0.043, *SE* = 0.016, *p* = 0.009]. This association intensified with reward history, as indicated by the positive interaction between neural set size marker and reward history [*β* = 0.026, *SE* = 0.013, *p* = 0.035], suggesting a positive influence of RL on WM management.

### Determinants of neural RPE markers

A core tenet of RL is that RPEs are computed as a difference in experienced vs expected outcome, and hence neural markers of RPE should be larger if preceded in that same trial by lower neural RL markers (i.e., lower expectations). Indeed, higher neural RPE markers were associated with lower preceding neural RL markers [*β* = -0.03, *SE* = 0.01, *p* < 0.001], representing reduced surprise during outcomes, and consistent with past findings.^27^ This effect held even when including the RPE from the RLWM model as an additional predictor, demonstrating that the neural RL markers reflecting subjective trial-wise expectations over and above what could be gleaned from the fitted RPEs based on behavior only (Suppl. Table S7)

As expected from RLWM principles, neural RPE markers also evolved across trials, decreasing with reward history but increasing under higher WM demand (higher set size and/or longer delays; Fig. 2F). Hence, neural RPEs declined as learned associations manifested, but remained more pronounced under higher WM demand (Fig. 2G) consistent with the cooperative RLWM model and past findings.^14,15,18^ In particular, we estimated a linear mixed-effect regression with neural RPE markers as dependent variable and predictors including set size, neural set size marker, and their interaction (Suppl. Table S8). This showed that neural RPE markers were pronounced under higher set size [*β* = 0.066, *SE* = 0.004, *p* < 0.001], highlighting the expected cooperative contribution of WM for RL-driven learning.^15^

The results discussed in Part I support RLWM model predictions and are consistent with previous work showing interactions between WM and RL in learning curves and in trial-by-trial neural indices of RL, WM and their interactions. Higher experienced WM demand (indexed by neural set size marker) increased the reliance on RL components by enhancing RPE signals.

### Part II: Findings Across Clinical Groups

Suppl. Fig. S2 visualizes the group-specific learning curves, demonstrating incremental learning in all groups with subtle differences. Overall, SCZ and BP had significantly lower accuracy than CTRL (Suppl. Fig. S3), while MDD and BP showed larger accuracy decrements as WM demand increased compared to CTRL (Fig. 3A). We will next show that learning impairments observed between clinical groups can be attributed to distinct neurocomputational sources. These sources became apparent in the model-based EEG analyses, as discussed in the following subsections, and in the comparisons of model parameters between groups, detailed in the final subsection.

**Figure 3.**
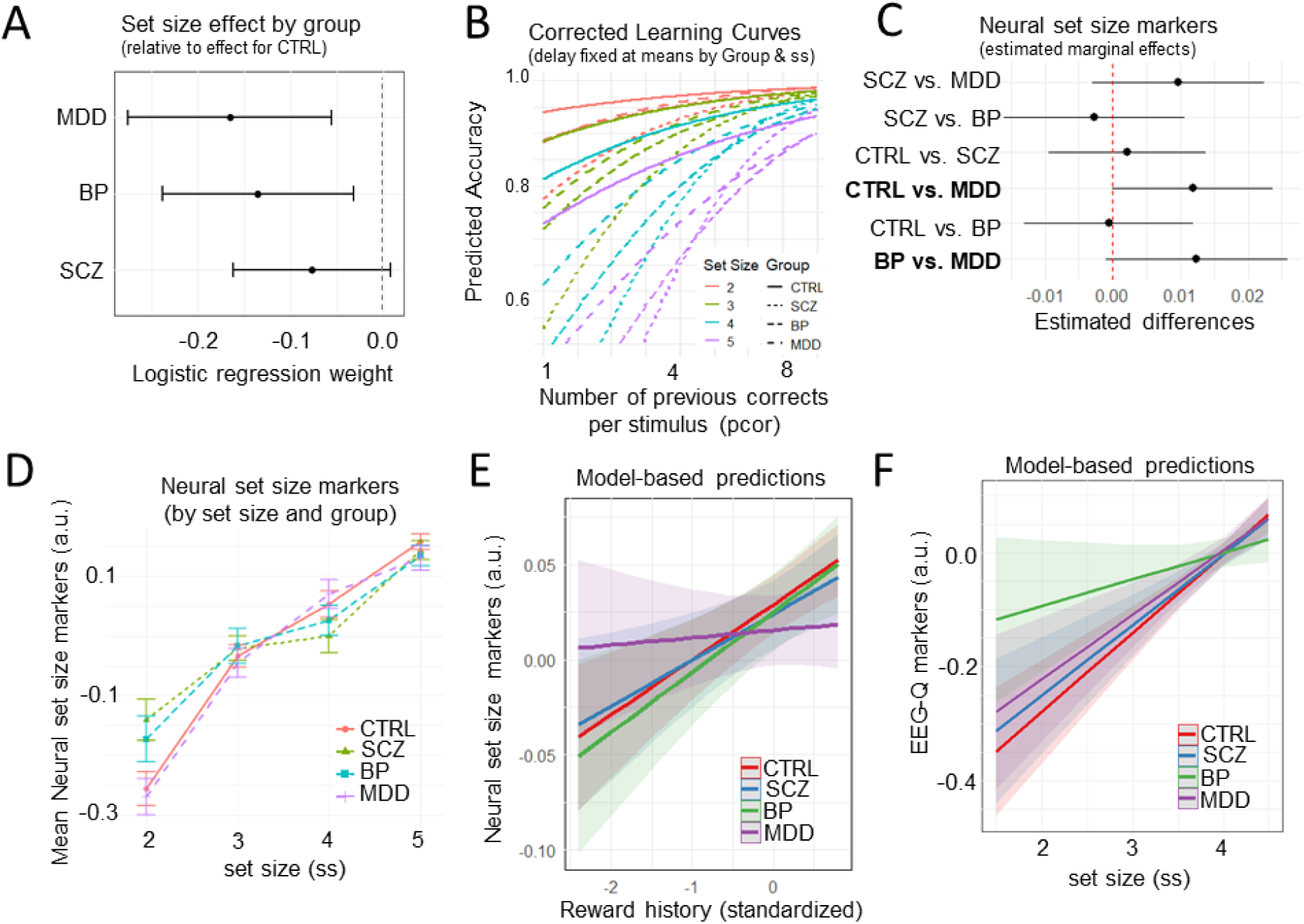
Depression (MDD) and Bipolar (BP) show similar disruptions with increasing set size but related to distinct model-based neural markers. **A.** Logistic regression with accuracy as dependent variable and predictors including delay, set size, reward history, group and their interactions. Dots represent point estimates and lines represent 95%-CI. This analysis showed larger set size effects for MDD and BP compared to CTRL. See Suppl. Table S10 for regression output and Suppl. Figure S4 for delay and reward history effects. **B.** Graphical illustration of the pronounced set size effect specific to MDD, BP, and SCZ with corrected learning curves by fixing delay at mean level within each set size and group. We also estimated a regression with accuracy as dependent variable and predictors including group, set size, delay, reward history, and their interactions. Compared to CTRL, BP and MDD had more trouble maintaining accuracy as set size increased across blocks. See Suppl. Figure S5 for plots by group. **C.** Pairwise group differences in coefficient weights from linear mixed-effect regression with neural set size marker as dependent variable and group as covariate. Dots represent point estimates and lines represent 95%-CI. See Suppl. Figure S6 for group comparison across all neural markers. **D.** Neural set size markers (arbitrary units) show parametric increases with set size (indexing higher WM demand), with weaker increases in BP and SCZ. Group means were calculated by first averaging within group and set size and then by group. Dots represent means and vertical bars represent SEMs. **E.** Model-based predictive plot of reward history on neural set size markers (arbitrary units) by group. Overall, participants showed increased engagement of neural indices of WM with reward history, this effect was blunted in MDD. Linear mixed-effect regression model included neural set size marker as dependent variable and predictors: set size, reward history, and group. See Suppl. Table S11 for regression output. **F.** Model-based predictive plot of set size effect on neural RL marker (arbitrary units) by group. Neural RL markers served as the dependent variable, while group, WM components (set size, neural set size markers), reward history, and their interactions served as predictors. See Suppl. Table S12 for regression output.

### Similar WM deficits in BP and MDD at behavioral levels

We first focus on how groups differed in their sensitivity to WM demand. Compared to CTRL, BP and MDD had more trouble maintaining accuracy as set size increased across blocks (Fig. 3A-B); see figure caption for regression model specification. The impact of set size on accuracy was moderated by the neural set size marker. Specifically, once neural set size was added as a predictor, the greater set size effects of MDD on accuracy rendered insignificant; a moderation that was not observed for CTRL or BP (Suppl. Table S9). Therefore, we next focused on group differences in neural set size markers.

### Distinct neural mechanisms contributing to WM deficits in BP and MDD

Although BP and MDD showed similar behavioral effects of set size, group comparisons in the model-based EEG markers suggested that these were due to distinct underlying mechanisms. MDD had significantly decreased neural set size markers compared to BP and CTRL, which did not differ from each other (Fig. 3C). Whether the reduced neural set size markers are due to less WM recruitment or deficits in WM management is not immediately evident. However, our computational analyses below suggest that reductions in neural set size markers in MDD were due to deficits in WM management. This is because MDD and CTRL groups displayed similar rises in neural set size markers with increased set size (Fig. 3D), suggesting comparable WM activation in response to higher WM demand. Moreover, we showed in the previous section that increased neural set size markers were associated with improved performance (i.e., higher accuracy).

### MDD is characterized by inefficient WM management

In part I, we showed that neural set size markers increased with reward history, putatively reflecting RL processes that recruit WM. Fig. 3E shows that this association was significantly weaker for MDD [*β* = -0.026, *SE* = 0.012, *p* = 0.003] compared to CTRL [*β* = 0.029, *SE* = 0.007, *p* < 0.001] who did not differ from SCZ [*β* = -0.003, *SE* = 0.012, *p* = 0.760] or BP [*β* = -0.002, *SE* = 0.012, *p* = 0.867]; see Suppl. Table S13. This result suggests deficits in the support of RL (reward history) on WM management (indexed by neural set size markers). Moreover, we did not find evidence for the alternative explanation of deficient WM recruitment in response to higher WM demand (Fig. 3D). Specifically, increases in set size were related to larger neural set size markers in CTRL [*β* = 0.134, *SE* = 0.007, *p* < 0.001] with no significant difference in MDD [*β* = -0.007, *SE* = 0.012, *p* = 0.532]. These findings distinguish MDD from BP and SCZ groups, who both showed deficits in WM recruitment rather than WM management as detailed next.

### BP and SCZ are characterized by reduced WM recruitment

BP and SCZ exhibited significantly smaller increases in neural set size markers under higher WM demand (increased set sizes) compared to MDD and CTRL [CTRL: *β* = 0.133, *SE* = 0.010, *p* < 0.001, MDD: *β* = -0.007, *SE* = 0.016, *p* = 0.661, SCZ: *β* = -0.039, *SE* = 0.016, *p* = 0.014, BP: *β* = -0.036, *SE* = 0.017, *p* = 0.037]; see Suppl. Table S14. This suggests potential deficits in WM recruitment, further supported by the observation that both groups experienced higher WM load at lower demands as indicated by their already elevated neural set size markers in blocks with lower set sizes (Fig. 3E).^10^

### BP is characterized by lower neural RL markers

Beyond the blunted WM recruitment, BP also showed significantly blunted neural RL markers compared to CTRL and MDD (Fig. 4A). The tendency to recruit neural RL with increasing set size (when RL is more useful) was significantly more pronounced in CTRL [*β* = 0.069, *SE* = 0.011, *p* < 0.001] than in BP [*β* = -0.046, *SE* = 0.019, *p* = 0.015] as shown in Fig. 3F. SCZ and MDD did not differ from CTRL [SCZ: *β* = -0.007, *SE* = 0.017, *p* = 0.673, MDD: *β* = -0.012, *SE* = 0.018, *p* = 0.486]. Moreover, the previously established negative association between neural set size markers and neural RL markers was significantly more pronounced in BP [*β* = -0.034, *SE* = 0.012, *p* = 0.004] and SCZ [*β* = -0.031, *SE* = 0.011, *p* = 0.004] as compared to CTRL [*β* = -0.314, *SE* = 0.007, *p* < 0.001]. MDD did not differ significantly from CTRL [*β* = -0.002, *SE* = 0.011, *p* = 0.086]; see Suppl. Table S12. These findings indicate that BP was characterized by deficits in both WM and RL recruitment (particularly under higher WM demand). Conversely, SCZ displayed deficits primarily in WM recruitment (irrespective of WM demand), while MDD showed challenges in adaptive WM management rather than recruitment.

**Figure 4.**
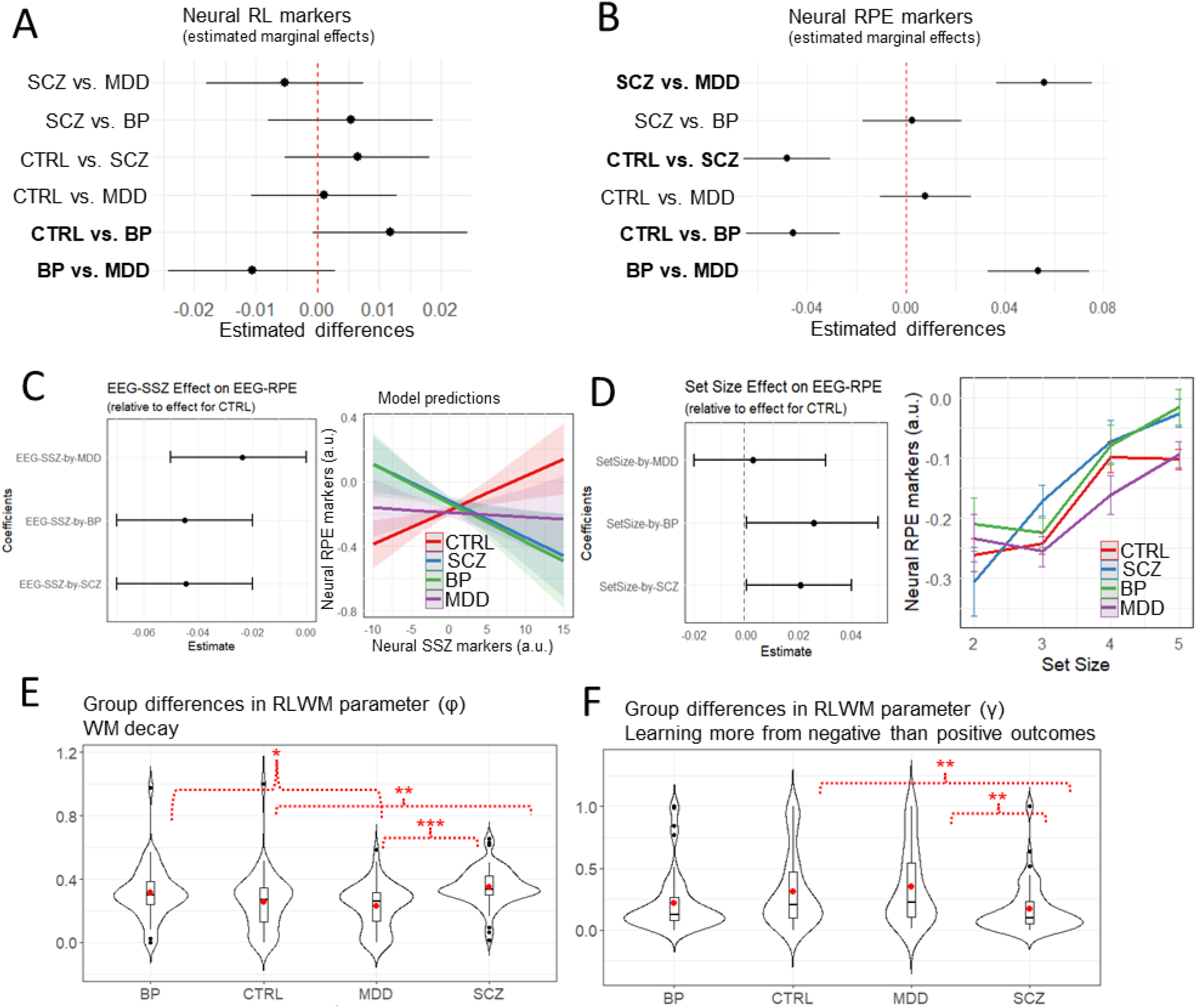
Bipolar (BP) and Schizophrenia (SCZ) show lower WM contributions to RL computations due to distinct underlying sources. **A.** Pairwise group differences in coefficient weights from linear mixed-effect regression with neural RL marker (of Q-values) as dependent variable and group as covariate. Dots represent point estimates and lines represent 95% confidence interval. **B.** Pairwise group differences in coefficient weights from linear mixed-effect regression with trial-based neural RPE marker as dependent variable and group as covariate. Dots represent point estimates and lines represent 95% confidence interval. **C.** Left plot: Coefficient from linear-mixed regression with neural RL-RPE marker as dependent variable and neural set size marker, set size, and group as predictors demonstrates grouping of BP and SCZ in terms of a diminished effect of WM onto RL computation. Right plot: Plotting neural RPE markers as a function of neural set size markers from the regression-based model shows the predicted positive association in CTRL (suggesting functional cooperative RL-WM interaction) but not in clinical groups. See Suppl. Table S15 for regression output. **D.** Left plot: Coefficient from linear-mixed regression with neural RL-RPE marker as dependent variable and task variables and group as covariates demonstrates larger neural set size effects for BP and SCZ compared to CTRL. Right plot: Plotting neural RPE markers across set size and by group shows greater activation for BP and SCZ under higher WM load (increased set size). Dots represent empirical data (averaged by subject and then by group). Vertical bars represent SEMs. See Suppl. Table S16 for regression output. **E.** Distribution of RLWM model parameter φ (WM decay) by group. Suppl. Fig. S7 provides group comparison across all parameters and Suppl. Fig. S8 shows distributions after outlier removal. The one-way ANOVA suggested a statistically significant difference in model parameter between the groups, *F*(3, 251) = 8.020, *p* < 0.001. To identify which groups differed significantly, a Tukey’s Honest Significant Difference (HSD) post-hoc test was performed. Significant results of the Tukey HSD test were: 1. Mean(MDD) minus Mean(BP) = -0.087, *95%-CI* = [-0.168, -0.006], *p-adjusted* = 0.030; 2. Mean(SCZ) minus Mean(CTRL) = 0.096, *95%-CI* = [0.031, 0.163], *p-adjusted* = 0.001; 3. Mean(SCZ) minus Mean(MDD) = 0.123, *95%-CI* = [0.123, 0.049], *p-adjusted* < 0.001. **F.** Distribution of model parameter (γ) by group. This parameter indexes learning more from negative than positive outcomes and is estimated by applying the RLWM computational model to accuracy as task performance measure. Suppl. Fig. S7 provides group comparison across all parameters and Suppl. Fig. S8 shows distributions after outlier removal. The one-way ANOVA suggested a statistically significant difference in model parameter between the groups, *F*(3, 251) = 6.058, *p* < 0.001. To identify which groups differed significantly, a Tukey’s Honest Significant Difference (HSD) post-hoc test was performed. Significant results of the Tukey HSD test were: 1. Mean(SCZ) minus Mean(CTRL) = -0.144, *95%-CI* = [-0.256, -0.031], *p-adjusted* = 0.006; 2. Mean(SCZ) minus Mean(MDD) = -0.178, *95%-CI* = [-0.305, -0.052], *p-adjusted* = 0.002.

### SCZ is characterized by weaker WM-RL cooperation under higher WM demand

As noted in the Introduction, if the WM and RL modules interact cooperatively, we would anticipate a positive relationship between neural markers of set size and RPEs (Fig. 1C). This positive correlation has been empirically confirmed in previous non-clinical studies.^27^ Here, we replicate this pattern in CTRL [*β* = 0.020, *SE* = 0.007, *p* = 0.009].^26,28,29^ However, Fig. 4C shows that this cooperative effect was reduced for all the clinical groups compared to CTRL [SCZ: *β* = -0.050, *SE* = 0.011, *p* = 0.001; BP: *β* = -0.044, *SE* = 0.012, *p* = 0.005; MDD: *β* = -0.023, *SE* = 0.012, *p* = 0.049], and even more so for SCZ under higher objective WM demand indexed by increased set size [*β* = -0.029, *SE* = 0.012, *p* = 0.014]; see Suppl. Table S15. These results suggest disruptions in the typically^27,28^ cooperative relationship between the RL and WM modules among the clinical groups. These effects were not simply due to blunted RPE signaling overall; indeed, when analyzed over all trials irrespective of neural WM markers, SCZ and BP exhibited significantly higher neural RPE markers compared to MDD and CTRL (Fig. 4B). Moreover, SCZ and BP had significantly higher increases in RPE under higher WM load as shown in Fig. 4D, as expected given that overall their accuracy was lower in these set sizes and thus positive outcomes are surprising. Thus, our finding of reduced neural RPEs with increasing neural markers of WM is particularly indicative of a disruption in cooperative interactions between these brain systems, putatively reflecting reduced impact of prefrontal cortex on RL signaling (see Discussion).

### Comparing model parameters between and within clinical groups

Comparing groups in RLWM model parameters allowed us to further characterize distinct sources of behavioral alterations in terms of latent computational mechanisms. Additionally, correlational analyses between neural set size marker (averaged by participant) and the subject-specific RLWM parameters also provided a better understanding of individual differences *within each group*.^3^ We complement the previous clinical findings with three additional insights.

First, BP and SCZ had significantly higher WM decay (φ) than MDD, which may explain why they showed deficits in WM recruitment and performance impairment already at lower WM demand as compared to MDD who showed deficits in WM management but not WM recruitment (Fig. 4E). Focusing on individual differences with between-subject correlational analyses, those with more WM decay (higher φ) showed lower neural RL markers (*r* = -0.424, *p* = 0.008). Given that lower neural RL markers were associated with lower accuracy, these results further support the previous conclusion that BP is characterized by deficits in both WM and RL recruitment.

Second, participants with MDD that relied more on WM than RL components (higher ρ) had lower neural set size markers (*r* = -0.293, *p* = 0.043). While greater reliance on WM might seem beneficial for performance, it can be detrimental especially when WM management is deficient, as observed in MDD. Indeed, those with higher WM reliance (ρ) performed worse under higher WM demand (increased set size; *β* = -7.77, *SE* = 0.92, *p* < 0.001) as shown in a regression with accuracy as dependent variable and predictors including set size, ρ, and their interactions.

Third, we established earlier that SCZ differed from BP in that they exhibited deficits only in WM recruitment, while BP showed deficits in both WM and RL recruitment. The RLWM parameter analyses revealed further that SCZ also demonstrated more perseveration, learning less from negative than positive outcomes (lower γ; see Fig. 4F).

In Supplementary Fig. S7, we compare the groups across all model parameters. Notably, we did not find group-level differences in undirected decision noise (ε). This does not contradict prior findings of increased behavioral stochasticity in clinical populations^27,28^ because our computational model includes multiple parameters (e.g., WM decay, learning rate, and WM-RL weighting) that can capture variability that is otherwise attributed to noise. This suggests that what appears as stochasticity in simpler models may reflect interpretable process-level dynamics when examined through a richer mechanistic lens. Moreover, our task’s explicit and deterministic structure and feedback design may have further mitigated random responding.

To evaluate the robustness of group differences in RLWM parameters, we conducted sensitivity analyses after removing outliers (detailed in the Supplement, section 1). These additional sensitivity analyses led to the exclusion of 56 participants (CTRL = 19, BP = 15, MDD = 5, SCZ = 17) who showed extreme values in one or more parameters. Fig. S8 shows that the overall pattern of group differences remained unchanged, with some effects appearing even stronger. The supplementary section 1 also shows that the repeated between-subject correlation analyses confirmed the results reported above. We also refer to Fig. S9 for additional recovery analyses of estimated model parameters.

## Discussion

The RLWM task has demonstrated good test-retest reliability.^10,12,15,19,20^ This study highlights its first application to characterize cognitive mechanisms across psychopathologies. Some groups showed similar behavioral impairments such as MDD and BP, who exhibited greater accuracy decrements under higher WM demand than SCZ who showed impairments already under lower WM demand. The model-based EEG analyses linked these behavioral impairments to distinct deficits in neural RL and WM components. Specifically, deficits in MDD were linked to inefficient WM management (i.e., reduced tendency to adapt neural RL with experience) paired with an over-reliance on WM processing. Conversely, BP showed impairments in both WM and RL recruitment and higher WM decay, exhibiting less cooperative RLWM interactions (irrespective of WM demand). SCZ expressed also impairments in WM recruitment and neglect of negative feedback, consistent with previous studies.^10,14^ The model-based EEG additionally revealed a reduction in cooperative RLWM interactions under higher WM demand. Our findings demonstrate the power of integrative model-based approaches for the identification of distinct cognitive and neural signatures across groups. Especially, the nuanced differences in learning deficits would have remained hidden without the model-based EEG analyses. This is important because MDD, BP, and SCZ have been commonly associated with learning impairments but the underlying mechanisms contributing to these impairments remained inconclusive thus far.^17,29^ Our study underscores the critical role of computational modeling and multi-level analysis in uncovering new insights.^3,9,17^

We also advance existing theories within the broader domain of computational cognitive neuroscience. Theoretical models of frontostriatal circuits have proposed that WM not only influences instrumental learning, but that RL reciprocally supports WM management.^25^ The presented findings provide empirical support for these theories, demonstrating that neural set size marker not only increased with subjective WM load under higher demand but also with reward history, suggesting improved WM management as RL processes alleviate the load on WM resources. To further elucidate the distinct roles of WM maintenance and management, future studies should explore these dynamics through novel manipulations that contrast simple decision-making processes with more complex ones.

Recent non-clinical studies suggest that RL and WM processes are not independent modules. Rather, behavioral, fMRI and EEG studies have suggested that WM influences RL.^14,15,18,19^ Thus, in principle, observations of diminished behavioral performance may partly reflect diminished influence of WM onto RL processes, which are posited to involve top-down prefrontal interactions with striatum. Indeed, previous studies have suggested that SCZ are less susceptible to top-down instructions linked to WM need to be held in prefrontal cortex (PFC) and typically inform striatal RL.^30^ The evidence presented here provides more direct support for reduced WM contributions to RL in SCZ, in that patients showed reduced influence of neural markers of WM onto neural markers of RPE in the same trial. This is important to consider given that past studies showed that behavioral impairments in RL tasks is more strongly tied to deficits in WM function, whereas RL-specific computations appear to be relatively intact in SCZ.^19,31^ Moreover, our finding of reduced neural WM markers in SCZ aligns with established endophenotypes involving deficits in attention, cognitive control, and abnormalities in sleep-related physiology, such as reduced sleep spindles and slow-wave activity.^32–35^ We view our approach as a bridge between mechanistic computational modeling and traditional endophenotypic frameworks: by estimating latent neurocognitive dynamics from behavior and EEG, future research can begin to map these task-derived markers onto broader biological traits, including sleep architecture and genetic risk profiles as well as their predictability of symptoms over time.

The interplay between WM and RL has been a key area of focus in cognitive studies, shedding light on how these systems collaboratively influence learning.^16,18,20^ Consistent with past findings, we found that higher experienced WM load (indexed by neural set size marker) increased the reliance on RL components by enhancing RPE signals.^15,18^ This aligns with neurocomputational models that emphasize dopamine’s role not only in RL mechanisms but also in modulating interactions with WM, affecting both learning and memory processes. Traditionally, dopamine has been primarily studied within the RL system, where phasic changes signal RPE in the striatum.^36–38^ Increasingly, studies are exploring how dopaminergic signaling related to RL also supports the manipulation and maintenance of WM content in the PFC.^25,39,40^ For instance, the prefrontal-basal ganglia neurocomputational model (PBWM^25^) highlights the intricate reciprocal interactions between WM and RL, emphasizing dopamine’s role across multiple cognitive processing systems. Beyond the computational gating mechanisms that integrate WM with RL processes in the basal ganglia, there is structural evidence that genetic factors affecting dopamine levels in the prefrontal cortex (PFC) influence the integrity of white matter.^20,41^ This integrity impacts the connections between WM-related brain components and the basal ganglia, which has been shown to corelate with variations in WM performance.^12,41^ Our findings underscore the importance of examining the dynamics and interactions between subsystems such as WM and RL in greater detail, and eventually developing a unified neurocomputational understanding. Although we did not examine spectral dynamics directly, the model-based EEG approach used here isolates neural markers of latent RL and WM computations without relying on predefined frequency bands or specific event-related components but rather focuses on extracting the latent mechanisms are simulated by the computational model. Prior work has shown that such voltage-based regressors yield robust signals of underlying computational processes, though future work may integrate both time- and frequency-domain analyses to further enrich mechanistic interpretations.

Recent work by Kirschner et al. (2024) has demonstrated the utility of EEG in identifying transdiagnostic learning impairments using ERP components such as the FRN and P3. While such approaches reveal important group-level associations, our study provides a complementary perspective focused on the mechanisms that give rise to neural dynamics and behavior. Specifically, our model-based EEG analyses dissociate reinforcement learning from working memory and link their trial-by-trial latent dynamics to observed choices in a way that is mechanistically interpretable. That is, grounded in specific cognitive computations like value updating and working memory engagement. In contrast, traditional ERP and time-frequency analyses often identify neural-behavior associations without clarifying which cognitive process the signal reflects and/or how it shapes behavior (see Discussion section in^3^ for a more detailed elaboration of this issue).

Examining how neurocognitive, motivational, and situational mechanisms interact at different timescales is crucial for building a more holistic understanding of symptom patterns across psychopathologies.^3–5,11^ Such a mechanistic understanding forms the basis for identifying and predicting the mechanisms to be targeted in interventions. This study sets the first groundwork by disentangling neurocomputational mechanisms contributing to clinical impairments within a specific context, namely in an abstract associative learning task. While this serves as an important starting point, it is essential to recognize that clinical symptoms often manifest and vary across different contexts. Future work should consider the interplay of neurocomputational sources of learning across multiple domains to fully understand symptom profiles.

Moving the field of computational psychiatry forward, future studies should address the following limitations. First, our study focused on differences and similarities across diagnostic categories. Future studies should examine symptom dimensions and severity within and across diagnoses to further dissect the mechanisms underlying learning impairments. This study serves as a foundational first step by demonstrating that even at the level of categorical diagnoses, often considered too broad for mechanistic insights, distinct neurocognitive processes can be measured using a computationally informed task design and model-based EEG analyses. By establishing that shared learning impairments at a behavioral level can arise from different latent mechanisms in MDD, BP, and SCZ, we lay a rigorous foundation for future dimensional and individualized approaches. Such layered modeling, moving from diagnostic groupings to transdiagnostic and symptom-specific signatures, will be critical for advancing precision psychiatry. Future research should also explore how latent neurocognitive profiles map onto individual differences in symptom expression and clinical outcomes to show the clinical utility of neurocomputational markers. Second, learning impairments likely reflect a confluence of cognitive, affective, and motivational processes, and their expression is shaped by environmental and task context. While this study focused on two central cognitive processes (RL and WM) in a cognitive task designed to dissociate between them, future research should extend this approach to incorporate broader functional and affective dimensions to create a richer assessment environment that allows distinct probing of not only cognitive but also affective and motivational processes. This is important because what appears as a learning deficit in a cognitive task may instead reflect motivational disengagement or impulsivity in another task with social elements or reward-specific manipulations. To disentangle these possibilities, a unified and calibrated battery of tasks is needed that systematically manipulates context along transdiagnostic dimensions (e.g., reward sensitivity, intolerance to uncertainty) while holding core parameters constant. Such designs will be critical for isolating context-general versus context-specific mechanisms and identifying which are most relevant to intervention targets. Future studies could also explore associations between model-derived mechanisms and performance on independent cognitive tasks. However, to ensure interpretability, it will be important that such tasks are specifically designed to dissociate the underlying computational processes, rather than conflating multiple mechanisms under a single construct, as is often the case with conventional working memory tasks. Third, we did not observe reduced perseveration in MDD, which is in contrast to prior research (using other paradigms than the RLWM task) albeit evidence remained mixed.^42–44^ It might be that the RLWM task did not elicit the necessary strong affective reactions to reveal dysfunctional reward processing. Studies suggest that not only perseveration but also emotional reactivity and intensity are distinctly shaped by affective traits associated with mood disorders and anxiety disorders that frequently co-occur with MDD.^45–47^ These findings underscore the need for neurocomputational assessments that probe both cognitive and affective processes for a more holistic understanding of mental health conditions. Emerging approaches, such as engineered game paradigms, show promise for expanding the scope of these assessments.^48,49^ Fourth, while individuals with SCZ, MDD, or BP often experience significant challenges in social interactions, these difficulties are underexplored in traditional neurocognitive testing.^3,50–53^ Recent advances suggest that game theoretical paradigms from experimental economics, coupled with neurocomputational modeling, offer powerful tools to investigate latent mechanisms in social and strategic learning contexts.^3,9,54,55^ While our primary goal was to explore latent computational and neural mechanisms underlying learning impairments, we acknowledge that group-level differences may be influenced by demographic variability. We deliberately attempted to match our clinical and control groups’ variables such as age, sex or race and parental education. Table 1 shows that we were successful in doing so for all but sex, which has known base rate differences across disorders. Future studies with demographically matched control groups will be important to more definitively dissociate diagnosis-related mechanisms from potential confounding factors. Our study allowed us to investigate learning impairments across psychopathologies, highlighting the need to broaden our approach beyond simple cognitive assessments. Developing multidimensional assessments allows researchers to explore the mechanisms of learning *and* their interaction with cognitive flexibility, enhancing our understanding of behavioral adaptability across different cognitive and social-cognitive contexts.^3,53^ Behavioral adaptability, an essential clinical characteristic, often goes underexplored, particularly in terms of its variability across different contexts and mental states.^1,53,56^ This variability is crucial because mental states are inherently dynamic, and cognitive processes such as WM and RL are not utilized in isolation. Instead, they are influenced by broad, fluctuating mental states which, if rigid, can impair error detection and adaptive responses. Our findings align with previous research indicating that the clinical groups displayed reduced adaptability to unexpected outcomes, driven by distinct neurobiological mechanisms. This underlines the importance of context-sensitive evaluations that can shed more light on how these disorders manifest in day-to-day life, ultimately guiding more effective, personalized interventions.

## Methods

### Participants

This study is part of the Cognitive Neuroscience Test Reliability and Computational Applications for Schizophrenia Consortium (CNTRaCS). All participants were recruited between January 2020 and June 2022 from local clinics and surrounding communities of five testing sites: the University of Maryland, the University of California, Davis, the University of Minnesota, the University of Chicago, and Washington University in St. Louis. All recruiting methods and experimental procedures were approved by a central IRB at Washington University. We provide demographics by group in Table 1.

Diagnostic status for all participant groups was established using the Structured Interview for the DSM-5 as described below. All patients were clinically stable with no medication changes in the previous month or anticipated in upcoming month. Additionally, participants with MDD met DSM-5 criteria for at least 2 depressive episodes, at least one of which occurred within the past three years. Participants with SCZ were diagnosed with either schizophrenia or schizoaffective disorder based on SCID interview described below. CTRL had no current psychiatric diagnosis, were not taking psychiatric medications, and reported no family history of psychosis. All participants were 18-63 years old, reported no history of neurologic injury, and were free from a substance use disorder in the three months prior to study enrollment.

After data preprocessing (described in subsequent sections), the analyses for this study included 223 participants (CTRL=71; SCZ=59; BP=45; MDD=48). 32 participants (CTRL=16, SCZ=8, MDD=6, BP=2) were excluded during data preprocessing procedures (see Methods: Participants). Specifically, 23 participants were removed from all the reported EEG analyses due to a high EEG artifact rate (>40% in one or more of the conditions in the stimulus or feedback locked data). 9 participants were removed because they had difficulties performing the task.

### Diagnostic Assessment

All participants completed a Structured Clinical Interview (SCID^57^) for DSM 5 with a trained researcher supervised by a doctoral-level clinician. Raters were trained with remote webinars in which rating scales and anchor points were discussed. Raters also completed and discussed a set of 6 training videos. Following this training, raters then worked to achieve consensus in their ratings with “gold standard” ratings that were supplied by experienced clinicians at the Maryland and St. Louis sites for at least 6 interviews. Consensus was defined as no more than 2 items with a difference of more than 1 rating point from the standard. To maintain inter-rater reliability over the course of the study, the St. Louis site recorded an interview to rate every 2-4 weeks, and all raters participated in remote meetings to resolve any discrepancies in ratings of this interview.

### RLWM Task

The RLWM task differentiates between RL and WM mechanisms by assessing the impact of distinct task manipulations on learning processes. Participants learn stimulus-response associations through trial-and-error and subsequent feedback in a learning phase that consists of multiple blocks of trials. Error responses are rewarded with zero points, while correct responses are probabilistically rewarded with either one or two points. The probabilistic variation in reward size is used to assess reward sensitivity in a subsequent reward retention test phase.^15,19^

The task is structured into multiple blocks of trials, with systematic manipulations both between and within blocks to parametrically adjust WM demand. Specifically, WM mechanisms are assessed by varying the set size between blocks and delay within blocks. Set size refers to the number of stimulus-response associations to be learned per block. Delay refers to the number of intervening trials since a stimulus was last correctly paired with a response.

The RLWM task differentiates between RL and WM mechanisms by assessing the impact of distinct task manipulations on learning processes. Specifically, RL-related mechanisms are quantified by the increase in accuracy with the accumulation of reward history, defined as the number of previous correct responses for a given stimulus-response association. Hence, RL refers to the incremental learning process that evolves over time throughout the task.

The task consisted of two phases: a learning phase and a subsequent reward retention test phase. For brevity, we report the main results of the reward retention phase in the Supplement, as they did not reveal clinically important findings.

### RLWM computational modeling

The RLWM model was employed to characterize the learning via two parallel processes - reinforcement learning (RL) and working memory (WM). The computational model was fit to the participants’ choice data. The RLWM model employed here is an adapted version from past studies.^15,18^

The RLWM model simulates behavior as a composition of contributions of RL and WM processes (Table 2). The fast but capacity-limited learning in the WM system contrasts with the slow but better retentive learning in the RL system. Both processes maintain a separate state-action value representation which is updated using temporal-difference style updates. However, the learning rate of the WM system α_*WM*_ is fixed at 1 to capture the fast, one-shot updating of the working memory which is capacity-limited and subject to decay over time. The WM capacity is denoted by an integer number of slots *C* and the decay is modulated by a decay parameter *φ*. The final action policy is derived from a weighted sum of RL and WM policies. The parameter *ρ* denotes the participant’s overall extent of reliance on the WM process (over the RL process) for making decisions.

#### Temporal-difference updates in RL and WM processes

The trial-wise learning updates for the RL and WM processes are given by the equations (1) and (2) in Table 2, where *s* refers to the current trial stimulus, *a* is the chosen action, *r*_*t*−1_ is the reward obtained at the current trial, *α* is RL learning rate and *γ* is the perseveration parameter. The perseveration parameter captures the neglect of negative feedback. Note that the reward values were binary coded as zero for the incorrect response or one for the correct response. As shown previously,^15^ the model fits did not improve when using a non-binary, 0-1-2 point system in the Q-learning updates.

#### WM decay

The model simulates WM decay by gradually reducing the WM-based Q-values toward their initial values of *1/n_a_*, with *n_a_* representing the number of possible actions. The rate of this decay is controlled by the parameter *φ*. Table 2, equation (3) denotes WM decay at every trial. We interpret decay effects as interferences because decay happens on every trial in which the subject sees a different stimulus than the one decayed. Moreover, the delay effects are larger with higher set size, indicating that it reflects interference.

#### Behavior policy

For each trial, the Q-values generated by the RL and WM processes are converted into choice action probabilities via a softmax function, shown in Table 2, equation (4), with a fixed inverse-temperature parameter, *β*. The *β* is consistently set to 100 across all subjects to avoid significant trade-offs with other model parameters. The derived policies, pol_RL_ and pol_WM_, are then integrated into a weighted mixture policy, pol_mix_, which depends on the (block-wise) degree of WM reliance, represented by ω. Table 2, equation (5) shows block-wise computation of *ω*. The *ρ* parameter is the participant’s baseline tendency to rely on WM across contexts (blocks), *C* parameter represents the participant’s WM capacity and ss is the set size or the number of unique stimuli encountered in each block. Table 2, equation (6) computes the mixture policy derived from pol_RL_ and pol_WM_.

#### Undirected noise

The final choice policy for the training phase, pol_final_, is computed as a weighted mixture of the model-derived choice policy (craven by RL and WM contributions) and a uniform random policy, which represents undirected noise or attentional lapses. This mixture is driven by the parameter *ε, which* estimates the probability of making a random choice, independent of learned or remembered values. Table 2, equation (7) formalizes this policy mathematically. Conceptually, *ε* captures latent decision noise or attentional disengagement and is commonly interpreted as a lapse rate in computational modeling.

#### Model fitting

The model is fit to the participant-level data using a constrained maximum likelihood estimation (MLE) procedure. We use Nelder-Mead optimization as implemented in *scipy.optimize*^58^ to get the MLE estimates. The parameter *C* was estimated as an integer value by looping MLE estimation over the integer parameter range [2,5] to get the best-fit combination of all parameters. Each estimate is obtained after running the optimization procedure with 40 different initializations (for each value of *C*) to increase the chance of finding the global optimum. All other parameters were constrained in range [0,1] except for the *α* parameter which was constrained to range [0,0.3] to avoid identifiability issues when α ∼ α_WM_. The previous RLWM modeling results^10,15,18,59^ have already shown that *α* << *α_WM_(1)* and *α* < 0.3. Note that following previous conventions of the RLWM task with the same model,^15,18,27^ α_WM_ is fixed at 1 to reflect the assumption that working memory updates deterministically upon feedback and has perfect recall. To maintain identifiability between the RL and WM systems, α must be kept sufficiently lower than 1. Otherwise, the systems become indistinguishable.

### EEG recording and processing

We used a Brain Products active electrode ActiCHamp system with 29 scalp electrodes and 3 electroocular electrodes for recording electrophysiological activity. A detailed description of our preprocessing pipeline is provided in previous publications.^60,61^ Although participants were recruited from different sites, we used a shared infrastructure and standardized testing protocols as well as analyses pipelines. Specifically, all behavioral and EEG data were collected using the same task implementation, hardware setup, EEG system and preprocessing pipeline. Testing procedures were harmonized across sites, including experimenter training and instructions to participants, and data quality was formally assessed on a regular basis to ensure similarity across sites.^60^

Specifically, preprocessing of EEG data was done in Matlab using the EEGLAB Toolbox^62^ and the ERPLAB plugin.^63^ We high-pass filtered the raw data (sampling rate: 500Hz; antialiasing filter: 130Hz) using a Butterworth filter (half-amplitude cutoff at 0.05Hz with a 12dB/octave slope) and screening for malfunctioning channels (i.e., channels with more than 1/3 unusable data). We then used Independent Component Analysis (ICA) to correct for eye blinks and horizontal eye movements artifacts. Channels identified as nonfunctional were excluded from the ICA and later interpolated using a spherical spline algorithm. About 67% of participants had no malfunctioning channels requiring interpolation (CTRL: 72.5%; SCZ: 60.34%; BP: 59.5%; MDD: 74.5%), while the rest had between 1 and 6 channels interpolated (average for CTRL: 2.56; SZ: 2.78; BP: 2.24; MDD: 2.54).

The continuous EEG was down-sampled to 125Hz, then was reduced to a selected window of - 100 to +700ms twice. Once it was epoched and baseline corrected from –100 to 0ms before the onset of the stimulus and second epoched and baseline corrected from –100 to 0ms before the onset of the feedback. The epoched data was subjected to an artifact detection algorithm (100µV voltage threshold with a moving window width of 200ms and a 100ms window step) followed by manual verification. Trials containing large artifacts were flagged and removed later at the univariate EEG analysis stage.

### Model-based EEG analyses

#### Data processing for univariate EEG analysis

To extract the neural correlates in the EEG signal of conditions of interest we employed a mass univariate approach.^18^ A multiple regression analysis was conducted for each participant, in which the EEG amplitude at each electrode site and time point was predicted by the conditions of interest: set-size (number of stimulus-response-outcome associations given in a block), model-derived RL expected value (denoted as Q), delay (number of trials since the same stimulus was presented and a correct response was given) and the interaction of these three regressors, while controlling for other factors like reaction time (log-transformed) and trial number within block. To account for remaining noise in the EEG data, the EEG signal (at each time point and electrode) was z-scored across all trials and so were all the predictors before they were entered to the robust multilinear regression analysis.^15,18^ The neural RL marker (of expected Q-values) and neural RPE marker were subsequently used as predictors in general mixed-effect regression analyses. The neural RL marker was calculated by averaging the values of the final 4 iterations of each stimulus to capture asymptotic performance. This was done for each stimulus-response association within each participant.

#### Corrected ERPs

To plot corrected ERPs, we computed the predicted voltage using the multiple-regression model described above while setting a single regressor to 0 (set size, delay, expected Q value, or reaction time); we subtracted this predicted voltage from the true voltage (for every electrode and time point within each trial), leaving only the fixed effect, the variance explained by that regressor, and the residual noise of the regression model. ERPs were computed as the average corrected voltage from all trials that belong to the same level of condition. Note that the array of expected Q values was divided to 4 quartiles and trials within each quartile were averaged for plotting ERPs.

#### Trial-by-trial similarity index of WM and RL

A multiple regression analysis was conducted for each participant, in which the EEG amplitude at each electrode site and time point was predicted by the conditions of interest (set size, delay, model-derived RL expected value, and their interactions). The delay predictor (the number of trials since the stimulus was presented and a correct response was made) used in the regression analyses was inverse transformed (-1/delay) to avoid the disproportion effect of very large but rare delays. We used the previously identified analysis method^15,18^ to identify spatiotemporal clusters (masks) of the three main predictors in the GLM (set-size, delay, and model-derived RL expected value). Specifically, we tested the significance of each time point at each electrode across participants against 0 using only trials with correct responses.

Importantly, as in previous research, we restricted our analysis to correct responses because we are interested in the underlying process of incremental learning. Focusing on correct responses then shows the dynamics of successful learning. This also avoids confounding effects due to positive or negative RPEs. In that sense, the RPE effects represent surprise and sensitivity to correct responses. This is because the actual percept a participant sees on the screen is the same on each trial. Hence, the only difference across trials is their learned expectations about reward. This is also why we expect it to be larger in high set size and systematically go down with experience. This approach ensures that all variance in the signal has to do with expectations, since that is the only thing that differs

For each marker we control what is presented to the subject. For instance, during choice, participants always look at one stimulus and prepare a response, but the neural WM markers reflect how many other stimuli they are learning about and how long it has been since they saw the current one; and the neural RL-Q marker reflects the reward expectation for that option based on their reward history and estimated learning rate.

#### Cluster statistics

We used cluster-mass correction by permutation testing with custom-written Matlab scripts for statistical inference. The spatiotemporal cluster mask is defined by the significant data points connected by temporal adjacency. The threshold for a t-test significance level for each time point is defined as P < (0.001). This was repeated 1000 times, generating a distribution of maximum cluster-mass statistics under the null hypothesis. Only clusters with a greater t-value sum than the maximum cluster mass obtained with 97.5% chance permutations were considered significant. Before using the mask to compute the similarity index in each trial we refined the mask and maintained only the highest effect size data points (10^th^ percentile) within each mask. We then assessed each trial’s neural similarity to the spatiotemporal mask of the condition by computing the dot product between the spatiotemporal voltage activity map of the individual trial and the spatiotemporal t-value map of the mask. This computation produced a trial-level similarity measure intended to assess the trial-wise experienced WM load and delay effects, as well as trial-wise RL contributions.

