## Supplemental Material for "Model-based EEG phenotyping uncovers distinct neurocomputational mechanisms underlying learning impairments across psychopathologies"

### 1. Additional Study Information

**Model fitting procedure.** We evaluated model fit at the individual level by computing the negative log-likelihood (NLL) of the fitted model for each subject, and derived the average likelihood per trial using the referenced formula. Subjects performing at or below chance (e.g., likelihood ~0.33) were flagged as potential non-compliant or poorly fit. None of the participants fell into this category (the highest value for one subject was found to be equal to 0.3).

**Reward retention test phase.** Stimuli that had been rewarded more often during learning were selected more often as having higher reward value in the test phase [ΔQ: $\beta = 1.11$, $SE = 0.09$, $p < 0.001$]. Moreover, performance in this test phase was enhanced when stimulus values were learned under higher than lower set size [mean(ss)-by-ΔQ: $\beta = 0.09$, $SE = 0.03$, $p = 0.001$] which is consistent with past findings.[2,3] This counterintuitive result is consistent with the EEG findings above and the WMRL interaction model, whereby larger RPEs during high WM load induce more robust RL computations that are expressed during the retention phase.

**Clinical differences in reward retention testing.** Data showed consistent performance at the test phase across clinical groups (p>.1) Note that we also controlled for response perseveration; no significant tendency was observed for repeating the same response used in the previous trial ($p > 0.20$).

**Sensitivity analyses of group differences in RLWM parameters after outlier removal.** To assess the robustness of the reported group differences in RLWM parameters (see Fig. S7), we conducted sensitivity analyses in which outliers were removed prior to re-analysis. Outliers were identified separately within each diagnostic group using the 1.5 x interquartile range (IQR) rule. Specifically, for each parameter, values falling below the first quartile minus 1.5×IQR or above the third quartile plus 1.5 x IQR were flagged as outliers. This non-parametric approach was chosen for its robustness to skewed distributions and group-specific variability. Participants with outlier values on any parameter were excluded from this secondary analysis. This procedure resulted in the exclusion of 56 participants (CTRL = 19, BP = 15, MDD = 5, SCZ = 17). The resulting parameter distributions are shown in Fig. S8. Hence, removal of outliers did not alter the pattern of group differences reported in Fig. S7 or in the final section of the Results in the main manuscript. These findings suggest that the reported effects are not driven by extreme values and are robust to outlier exclusion. We also repeated the between-subject correlation analyses reported in the last subsection of the Results, this time excluding participants identified as outliers based on model parameters. The findings remained robust: First, within the BP group, participants with greater working memory decay (higher $\varphi$) continued to show reduced neural reinforcement learning markers ($r = -0.48$, $p = 0.006$). Second, within the MDD group, participants who relied more on working memory than reinforcement learning processes (higher $\rho$) still exhibited reduced neural set size markers ($r = -0.35$, $p = 0.021$).

#### 2. Supplementary Figures

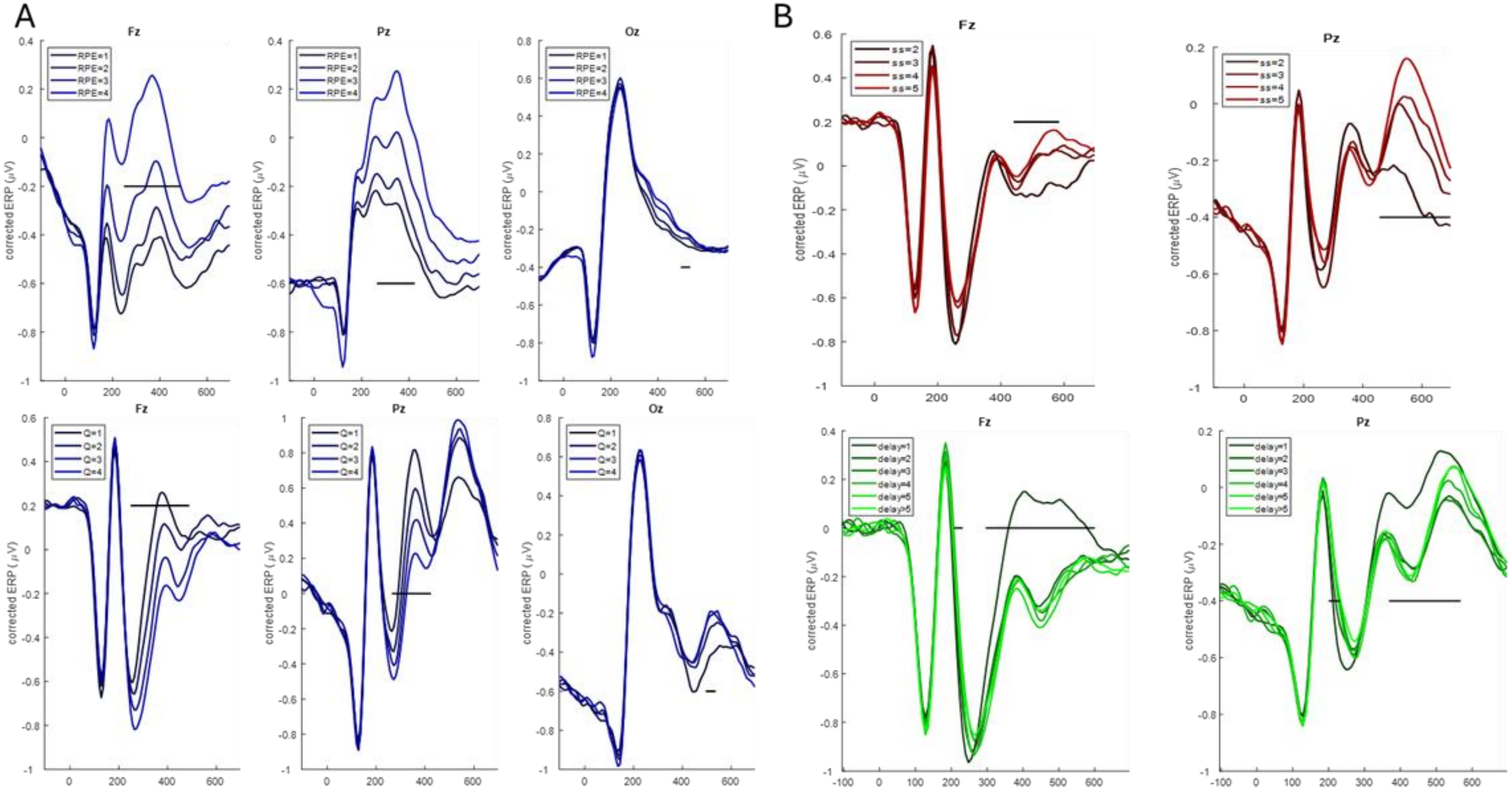


**Fig. S1. Event-related potentials (ERPs) by markers of reinforcement learning (RL) and working memory (WM).**

**(A)** ERP plots show the effect of the extracted RL markers, reward prediction errors (RPEs) and Q-values, derived from computational modeling and GLM (detailed in the Methods). Shown are the time course for four quartile values. Horizontal black lines reflect significant time points after permutation correction. **(B)** ERP plots show the effect of working memory markers (set size and delay) on the voltage of significant electrodes (Fz and Pz both at central lines). Horizontal black lines reflect significant time points after permutation correction.

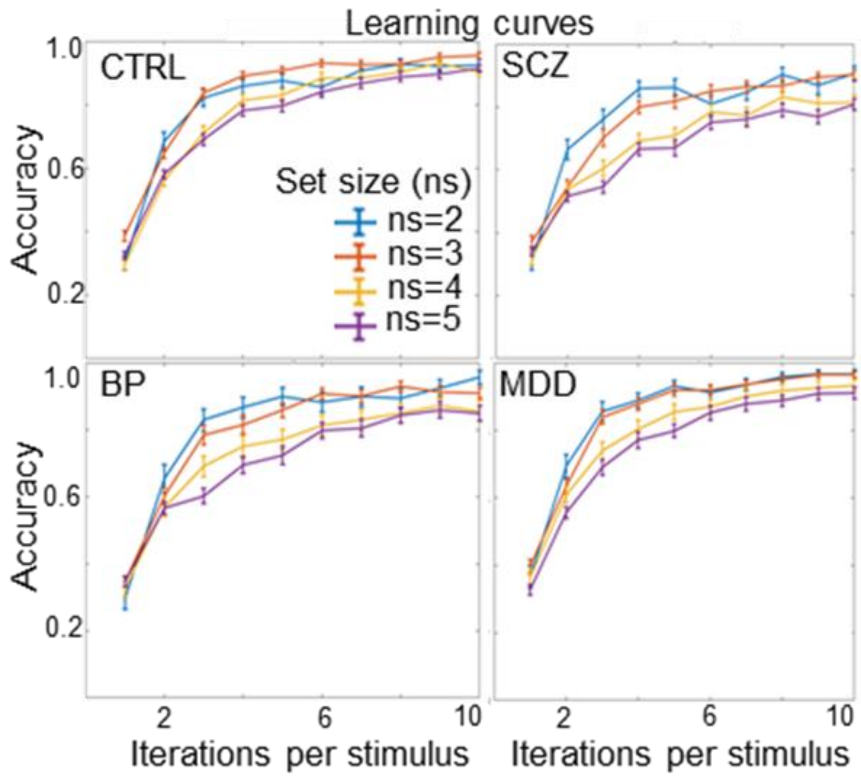


**Fig. S2. Learning curves by clinical group.**

Accuracy curves (averaged across trials, participants, and groups) by number of stimulus iterations show continuous learning over time that varies with set size in all clinical groups. Vertical bars represent within-subject SEs. CTRL refers to participants without any mental health diagnoses; SCZ refers to participants diagnosed with schizophrenia; BP refers to participants diagnosed with bipolar disorder; MDD refers to participants diagnosed with major depressive disorders.

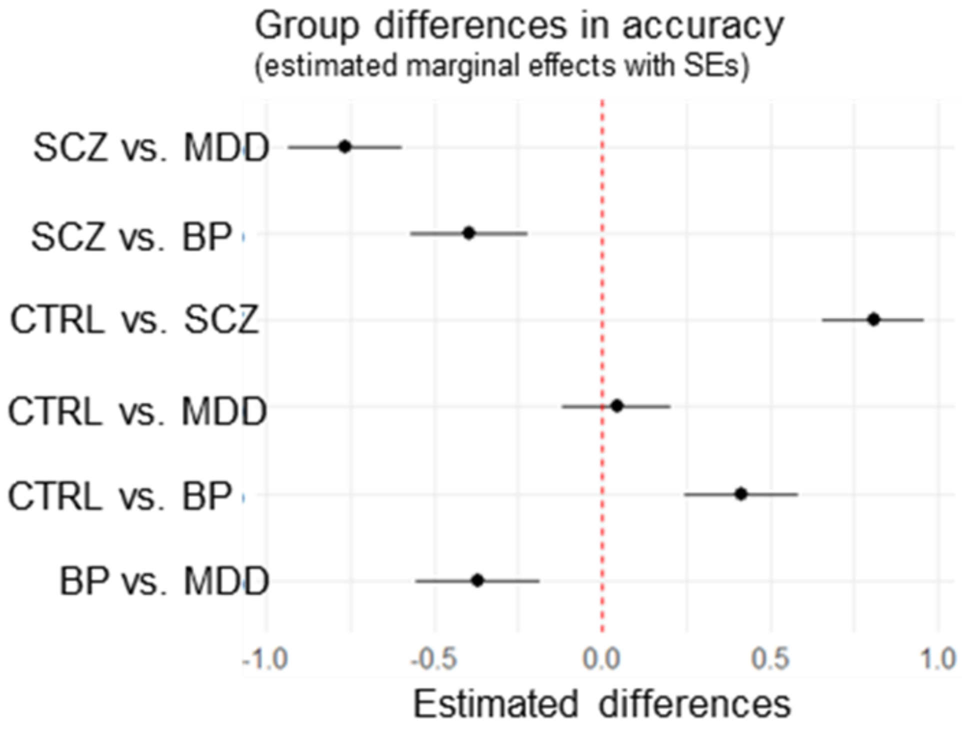


**Fig. S3. Group-wise comparison in overall accuracy (across all trials).**

Results from regression-based models with accuracy (across all trials) as dependent variable and group as independent variable. Points refer to means and horizontal lines refer to estimated standard errors. The CTRL and MDD group had an overall higher accuracy than the SCZ and BP groups. The CTRL and MDD group did not differ in overall accuracy. The BP group had a higher accuracy than the SCZ group. CTRL refers to participants without any mental health diagnoses; SCZ refers to participants diagnosed with schizophrenia; BP refers to participants diagnosed with bipolar disorder; MDD refers to participants diagnosed with major depressive disorders.

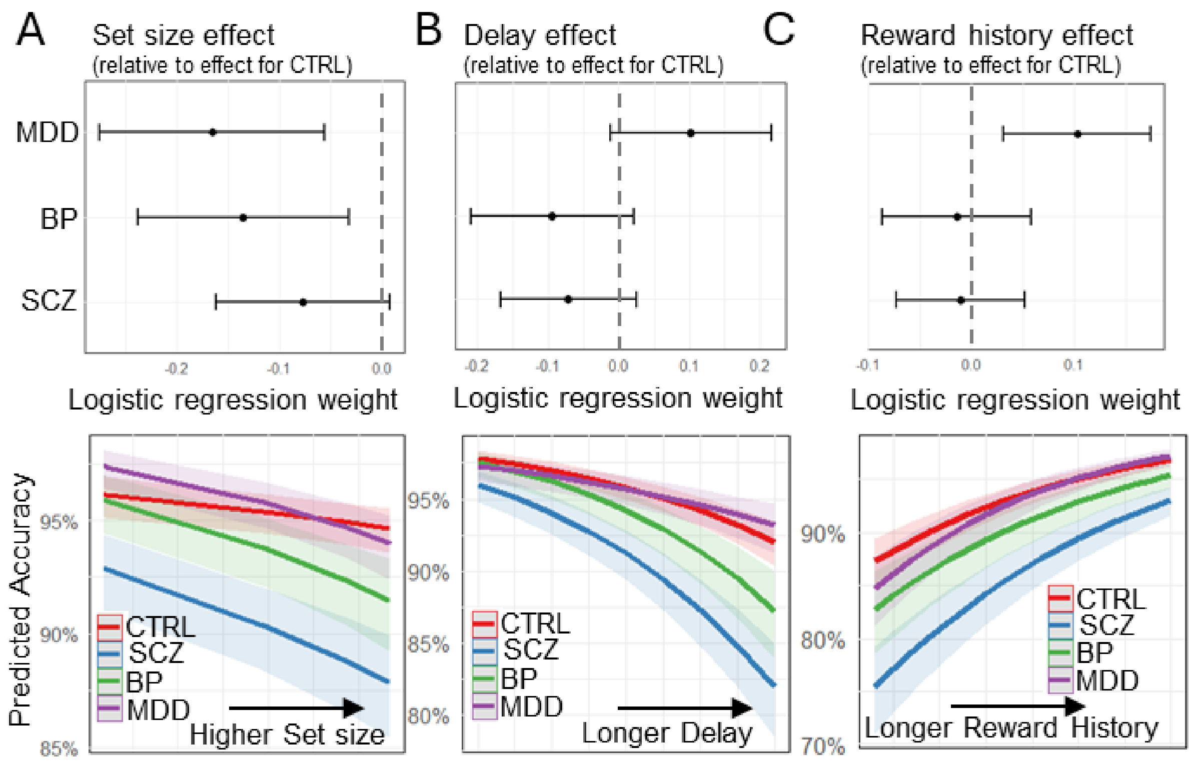


**Fig. S4. Group differences in set size, delay, and reward history effects.**
Results from logistic regression with accuracy as dependent variable and predictors including delay, set size, reward history, group and their interactions. Dots represent point estimates and lines represent 95%-CI. This analysis showed larger set size effects for MDD and BP compared to CTRL **(A)** as well as smaller reward history effects for MDD compared to CTRL **(C)**. We did not find significant group differences in delay effects **(B)**. See Suppl. Table S10 for regression output. CTRL refers to participants without any mental health diagnoses; SCZ refers to participants diagnosed with schizophrenia; BP refers to participants diagnosed with bipolar disorder; MDD refers to participants diagnosed with major depressive disorders.

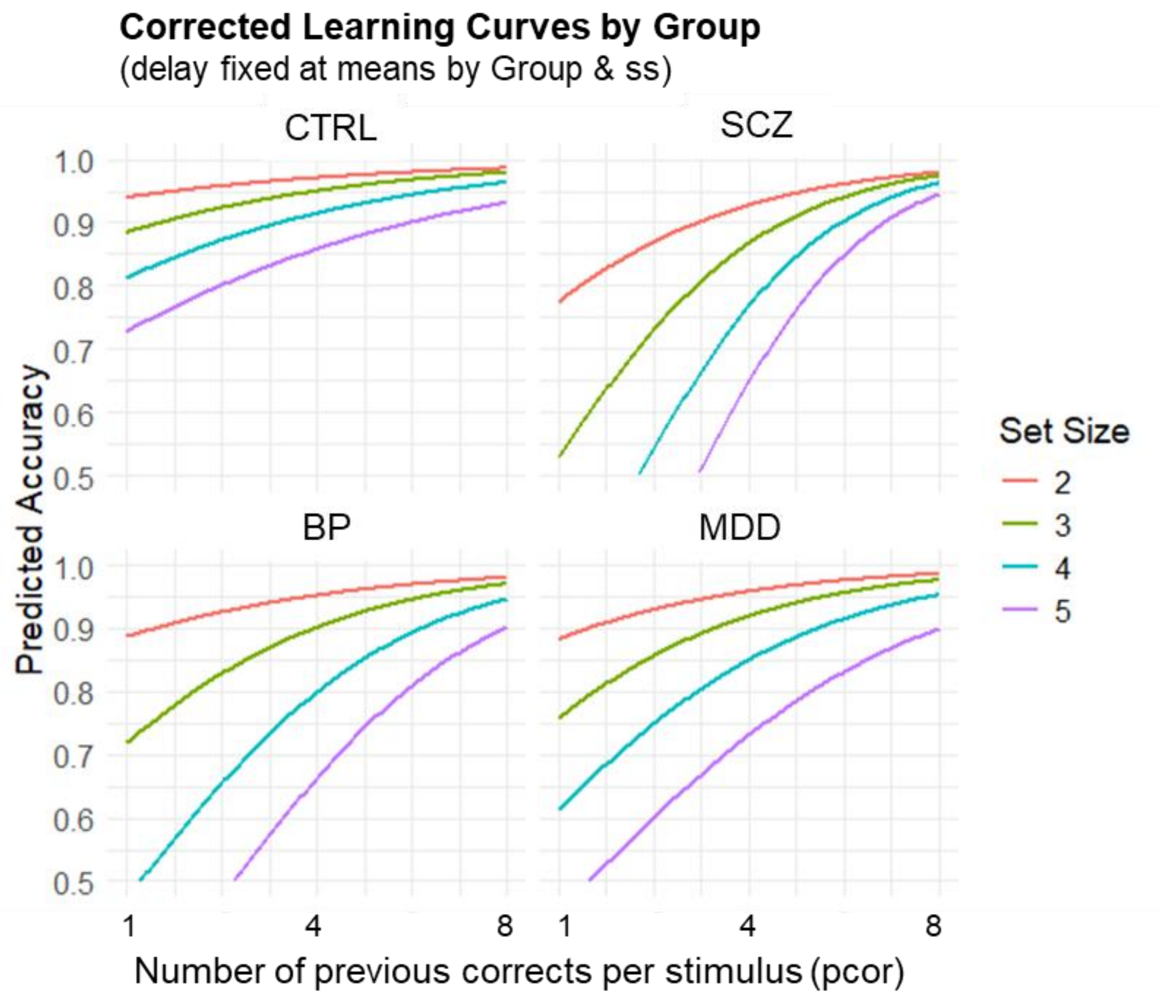


**Fig. S5. Delay-corrected learning curve by group.**

Graphical illustration of the pronounced set size effect specific to MDD, BP, and SCZ with corrected learning curves by fixing delay at mean level within each set size and group. We also estimated a regression with accuracy as dependent variable and predictors including group, set size, delay, reward history, and their interactions. Compared to CTRL, BP and MDD had more trouble maintaining accuracy as set size increased across blocks. CTRL refers to participants without any mental health diagnoses; SCZ refers to participants diagnosed with schizophrenia; BP refers to participants diagnosed with bipolar disorder; MDD refers to participants diagnosed with major depressive disorders.

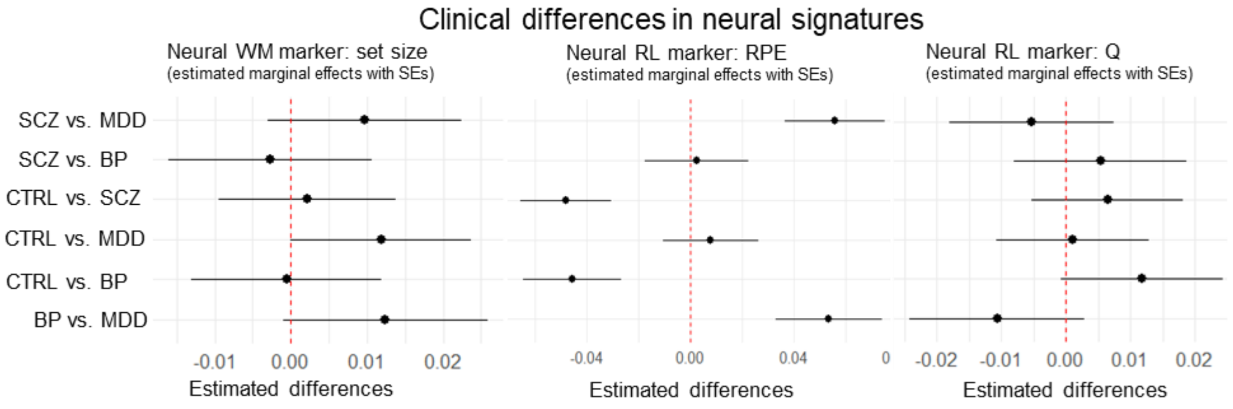


**Fig. S6. Group differences in neural markers of working memory (WM) and reinforcement learning (RL).**

Results from regression-based models with neural WM markers (set size) and reinforcement learning markers (RPEs, Q-values) as dependent variable and group as independent variable. We do not show neural markers of delay effects since we did not find significant group differences (see Supplementary Fig. S4B). Points refer to means and horizontal lines refer to estimated standard errors. **(A)** The CTRL group had overall higher neural set size markers than the MDD group. **(B)** The SCZ and BP groups had higher neural RPE markers than the CTRL and MDD groups. **(C)** The CTRL group had slightly higher neural Q markers than the CTRL group. CTRL refers to participants without any mental health diagnoses; SCZ refers to participants diagnosed with schizophrenia; BP refers to participants diagnosed with bipolar disorder; MDD refers to participants diagnosed with major depressive disorders.

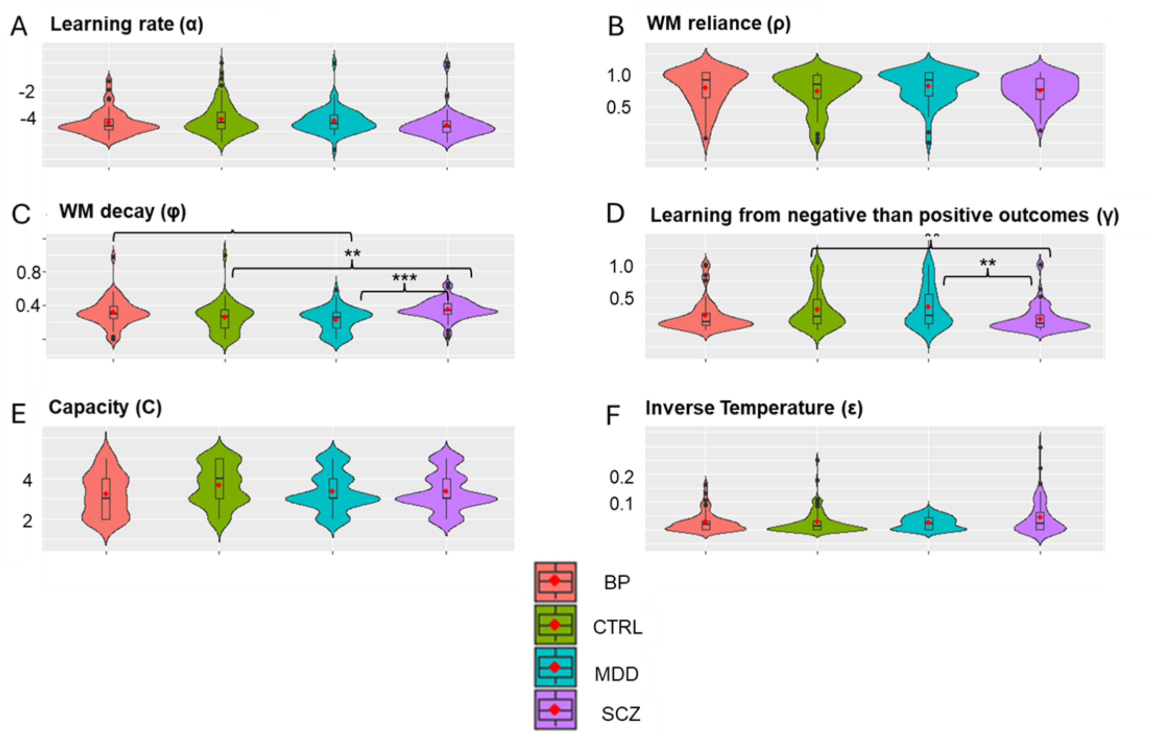


**Fig. S7. Group differences in computational RLWM model parameters.**

Shown are distribution of RLWM parameters with the red dots representing group means and the vertical boxplots representing the median (thick horizontal lines) and interquartile ranges. CTRL refers to participants without any mental health diagnoses; SCZ refers to participants diagnosed with schizophrenia; BP refers to participants diagnosed with bipolar disorder; MDD refers to participants diagnosed with major depressive disorders. Note that the parameter β is fixed at a value of 100 to prevent significant trade-offs with other parameters during model fitting. For details, see Method section in the main manuscript. **(A-B)** Shown are the distribution of RLWM model parameter by group for **(A)** learning rate (α) and **(B)** WM reliance (ρ) for which we did not find any significant group differences. **(C)** Distribution of RLWM model parameter φ (WM decay) by group. The one-way ANOVA suggested a statistically significant difference in model parameter between the groups, $F(3, 251) = 8.020$, $p < 0.001$. To identify which groups differed significantly, a Tukey's Honest Significant Difference (HSD) post-hoc test was performed. Significant results of the Tukey HSD test were: 1. Mean(MDD) minus Mean(BP) = -0.087, *95%-CI* = [-0.168, -0.006], *p-adjusted* = 0.030; 2. Mean(SCZ) minus Mean(CTRL) = 0.096, *95%-CI* = [0.031, 0.163], *p-adjusted* = 0.001; 3. Mean(SCZ) minus Mean(MDD) = 0.123, *95%-CI* = [0.123, 0.049], *p-adjusted* < 0.001. **(D)** Distribution of RLWM model parameter (γ) by group. This parameter indexes learning more from negative than positive outcomes and is estimated by applying the RLWM computational model to accuracy as task performance measure. The one-way ANOVA suggested a statistically significant difference in model parameter between the groups, $F(3, 251) = 6.058$, $p < 0.001$. To identify which groups differed significantly, a Tukey's Honest Significant Difference (HSD) post-hoc test was performed. Significant results of the Tukey HSD test were: 1. Mean(SCZ) minus Mean(CTRL) = -0.144, *95%-CI* = [-0.256, -0.031], *p-adjusted* = 0.006; 2. Mean(SCZ) minus Mean(MDD) = -0.178, *95%-CI* = [-0.305, -0.052], *p-adjusted* = 0.002. **(E-F)** Shown are the distribution of RLWM model parameter by group for **(E)** capacity (C) and **(F)** inverse temperature (ε) for which we did not find any significant group differences.

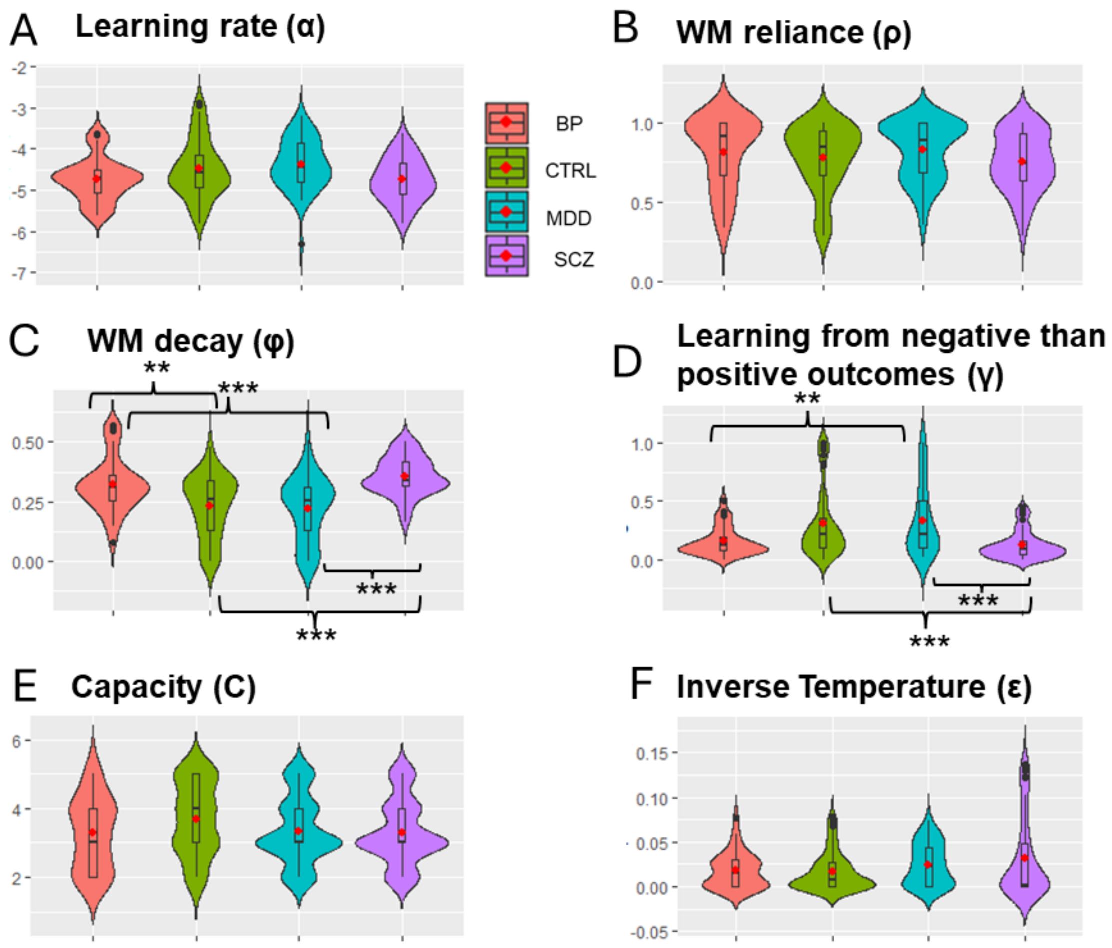


**Fig. S8. Group differences in RLWM model parameters after outlier removal.**

Shown are distribution of RLWM parameters with the red dots representing group means and the vertical boxplots representing the median (thick horizontal lines) and interquartile ranges. CTRL refers to participants without any mental health diagnoses; SCZ refers to participants diagnosed with schizophrenia; BP refers to participants diagnosed with bipolar disorder; MDD refers to participants diagnosed with major depressive disorders. Note that the parameter β is fixed at a value of 100 to prevent significant trade-offs with other parameters during model fitting. For details, see Method section in the main manuscript. **(A-B)** Shown are the distribution of RLWM model parameter by group for **(A)** learning rate (α) and **(B)** WM reliance (ρ) for which we did not find any significant group differences. **(C)** Distribution of RLWM model parameter φ (WM decay) by group. The one-way ANOVA suggested a statistically significant difference in model parameter between the groups, $F(3, 195) = 16.22$, $p < 0.001$. To identify which groups differed significantly, a Tukey's Honest Significant Difference (HSD) post-hoc test was performed. Significant results of the Tukey HSD test were: 1. Mean(CTRL) minus Mean(BP) = -0.089, *95%-CI* = [-0.154, -0.023], *p-adjusted* = 0.003; 2. Mean(MDD) minus Mean(BP) = -0.101, *95%-CI* = [-0.171, -0.032], *p-adjusted* = 0.001; 3. Mean(SCZ) minus Mean(CTRL) = 0.124, *95%-CI* = [0.067, 0.181], *p-adjusted* < 0.001; 4. Mean(SCZ) minus Mean(MDD) = 0.137, *95%-CI* = [0.075, 0.198], *p-adjusted* < 0.001. **(D)** Distribution of RLWM model parameter (γ) by group. This parameter indexes learning more from negative than positive outcomes and is estimated by applying the RLWM computational model to accuracy as task performance measure. The one-way ANOVA suggested a statistically significant difference in model parameter between the groups, $F(3, 195) = 9.515$, $p < 0.001$. To identify which groups differed significantly, a Tukey's Honest Significant Difference (HSD) post-hoc test was performed. Significant results of the Tukey HSD test were: 1. Mean(MDD) minus Mean(BP) = 0.181, *95%-CI* = [0.037, 0.324], *p-adjusted* = 0.007; 2. Mean(SCZ) minus Mean(CTRL) = -0.187, *95%-CI* = [-0.304, -0.069], *p-adjusted* < 0.001; 3. Mean(SCZ) minus Mean(MDD) = -0.212, *95%-CI* = [-0.339, -0.086], *p-adjusted* < 0.001. **(E-F)** Shown are the distribution of RLWM model parameter by group for **(E)** capacity (C) and **(F)** inverse temperature (ε) for which we did not find any significant group differences.

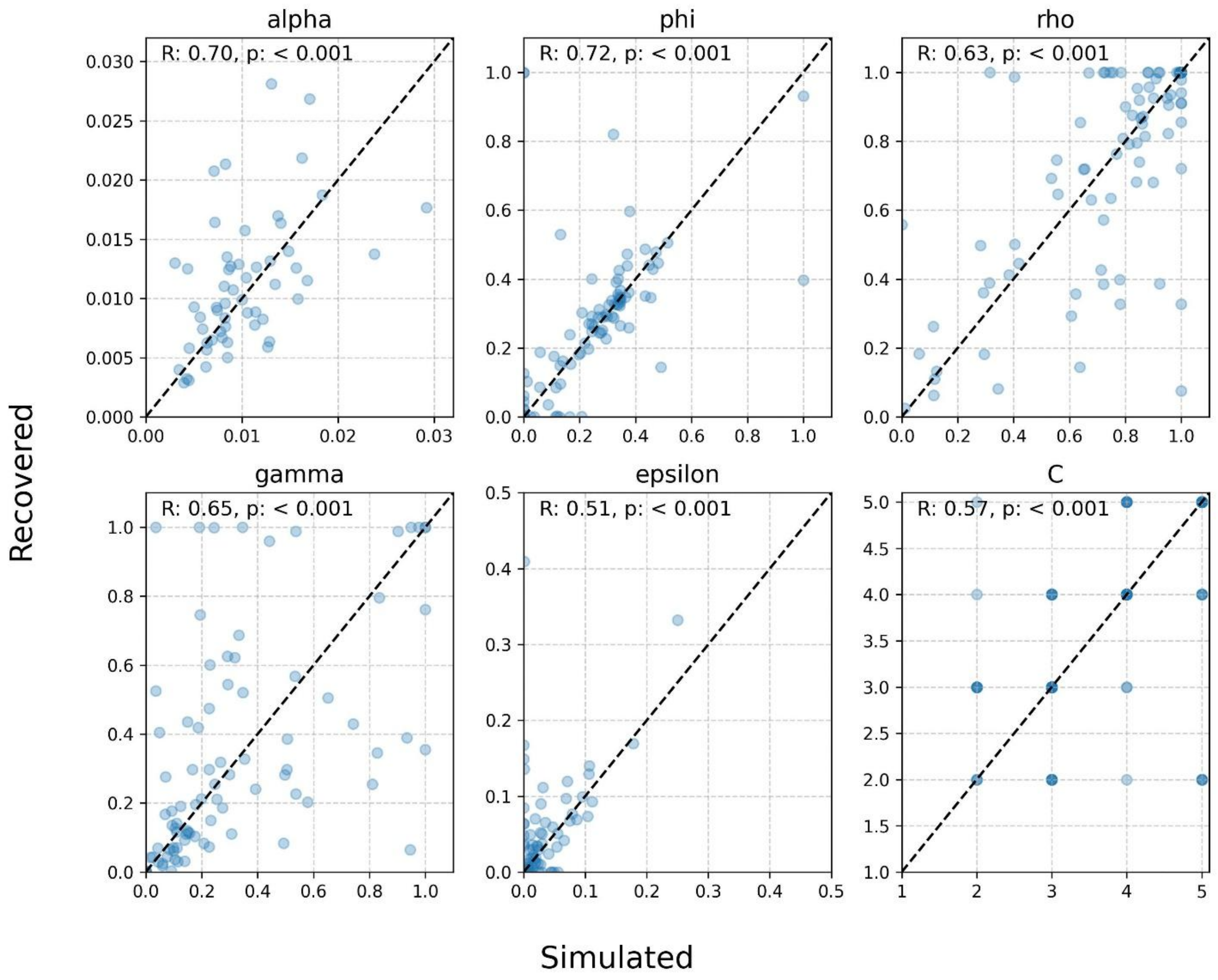


**Fig. S9. Parameter recovery based on simulated data.**

Results from simulations comparing input parameters (x-axis) and recovered parameters (y-axis). Overall, these plots show good parameter recovery (R refers to correlation coefficients).

### 3. Supplementary Tables

**Table S1. Regression output of model with accuracy as dependent variable.**

| **Dependent variable: Accuracy** | | | |
|---|---|---|---|
| *Predictors* | *Odds Ratios* | *CI* | *p* |
| (Intercept) | 16.69 | 14.52 – 19.17 | **<0.001** |
| Set Size | 0.75 | 0.70 – 0.80 | **<0.001** |
| Delay | 0.77 | 0.72 – 0.82 | **<0.001** |
| Pcor | 1.64 | 1.58 – 1.69 | **<0.001** |
| Set Size × Delay | 0.67 | 0.64 – 0.70 | **<0.001** |
| Set Size × Pcor | 1 | 0.95 – 1.04 | 0.843 |
| Delay × Pcor | 1.25 | 1.20 – 1.31 | **<0.001** |
| **Random Effects** | | | |
| $\sigma^2$ | 3.29 | | |
| $\tau_{00\ sbj}$ | 1.14 | | |
| $\tau_{11\ sbj.SetSize}$ | 0.12 | | |
| $\tau_{11\ sbj.Delay}$ | 0.09 | | |
| $\tau_{11\ sbj.Pcor}$ | 0.02 | | |
| $\tau_{11\ sbj.SetSize:Delay}$ | 0.02 | | |
| $\tau_{11\ sbj.SetSize:Pcor}$ | 0.02 | | |
| $\tau_{11\ sbj.Delay:Pcor}$ | 0.01 | | |
| $\rho_{01}$ | -0.1 | | |
| | 0.58 | | |
| | 0.35 | | |
| | -0.38 | | |
| | 0.08 | | |
| | 0.56 | | |
| N $_{sbj}$ | 255 | | |
| Observations | 68475 | | |
| Marginal $R^2$ | 0.154 | | |

Output from multi-linear mixed regression model fitted across all participants, complementing regression coefficient in Figure 2B of main manuscript. Regression model included accuracy as binary dependent variable, corrects (1) and errors (0), delay, and previous correct responses indexing reward history (pcor) as well as their two way interactions as independent variables. Shown are means and 95% confidence intervals of fixed effects as well as estimated random effects.

**Table S2. Regression output of model with EEG-Q as dependent variable.**

| **Dependent variable: EEG-Q** | | | |
|---|---|---|---|
| *Predictors* | *Estimates* | *CI* | *p* |
| (Intercept) | 0.05 | 0.05 – 0.06 | **<0.001** |
| Set Size | 0 | -0.01 – 0.01 | 0.776 |
| Delay | 0.05 | 0.04 – 0.05 | **<0.001** |
| Pcor | 0.11 | 0.10 – 0.12 | **<0.001** |
| Set Size × Delay | -0.01 | -0.01 – 0.00 | 0.191 |
| Set Size × Pcor | 0 | -0.01 – 0.01 | 0.947 |
| Delay × Pcor | 0.01 | -0.00 – 0.02 | 0.06 |
| **Random Effects** | | | |
| $\sigma^2$ | 0.98 | | |
| $\tau_{00\ sbj}$ | 0 | | |
| $N_{sbj}$ | 222 | | |
| Observations | 51794 | | |
| Marginal $R^2$ | 0.014 | | |

Output from multi-linear mixed regression model fitted across all participants, complementing regression coefficient in Figure 2D of main manuscript. Mixed-effect regressions included neural RL markers as dependent variable and predictors: set size, delay, reward history (pcor), and their interactions. Shown are means and 95% confidence intervals of fixed effects as well as estimated random effects.

**Table S3. Regression output of model with EEG-RPE as dependent variable.**

| **Dependent variable: EEG-RPE** | | | |
|---|---|---|---|
| *Predictors* | *Estimates* | *CI* | *p* |
| (Intercept) | -0.16 | -0.17 – -0.15 | **<0.001** |
| Set Size | 0.06 | 0.05 – 0.07 | **<0.001** |
| Delay | 0.03 | 0.02 – 0.04 | **<0.001** |
| Pcor | -0.17 | -0.18 – -0.16 | **<0.001** |
| Set Size × Delay | 0.04 | 0.03 – 0.04 | **<0.001** |
| Set Size × Pcor | -0.02 | -0.03 – -0.01 | **<0.001** |
| Delay × Pcor | -0.05 | -0.06 – -0.04 | **<0.001** |
| **Random Effects** | | | |
| $\sigma^2$ | 0.93 | | |
| $\tau_{00\ sbj}$ | 0.01 | | |
| $N_{sbj}$ | 222 | | |
| Observations | 51794 | | |
| Marginal $R^2$ | 0.039 | | |

Output from multi-linear mixed regression model fitted across all participants, complementing regression coefficient in Figure 2F of main manuscript. Mixed-effect regressions included neural reward prediction error (RPE) markers as dependent variable and predictors: set size, delay, reward history (pcor), and their interactions. Shown are means and 95% confidence intervals of fixed effects as well as estimated random effects.

**Table S4. Regression output of model with EEG-Q as dependent variable.**

| **Dependent variable: EEG-Q** | | | |
|---|---|---|---|
| *Predictors* | *Estimates* | *CI* | *p* |
| (Intercept) | 0 | -0.01 – 0.01 | 0.992 |
| Pcor | 0.12 | 0.11 – 0.13 | **<0.001** |
| Neural set size marker | -0.33 | -0.35 – -0.32 | **<0.001** |
| Set Size | 0.05 | 0.04 – 0.07 | **<0.001** |
| Pcor × Neural set size marker | 0.01 | 0.00 – 0.02 | **0.02** |
| Pcor × Set Size | 0 | -0.00 – 0.01 | 0.382 |
| Neural set size marker × Set Size | 0 | -0.01 – 0.01 | 0.515 |
| Pcor × Neural set size marker × Set Size | -0.01 | -0.01 – 0.00 | 0.198 |
| **Random Effects** | | | |
| $\sigma^2$ | 0.86 | | |
| $\tau_{00\ sbj}$ | 0 | | |
| $\tau_{11\ sbj.Pcor}$ | 0 | | |
| $\tau_{11\ sbj.NeuralSetSizeMarker}$ | 0.01 | | |
| $\tau_{11\ sbj.SetSize}$ | 0.01 | | |
| $\tau_{11\ sbj.Pcor\_by\_NeuralSetSizeMarker}$ | 0 | | |
| $\tau_{11\ sbj.Pcor\_by\_SetSize}$ | 0 | | |
| $\tau_{11\ sbj.SetSize\_by\_NeuralSetSizeMarker}$ | 0 | | |
| $\tau_{11\ sbj.Pcor\_by\_SetSize\_by\_NeuralSetSizeMarker}$ | 0 | | |
| $\rho_{01}$ | 0.2 | | |
| | 0.99 | | |
| | 0.19 | | |
| | 0.22 | | |
| | 0.05 | | |
| | -0.13 | | |
| | -0.02 | | |
| $N_{sbj}$ | 222 | | |
| Observations | 51794 | | |
| Marginal $R^2$ | 0.125 | | |

Output from multi-linear mixed regression model fitted across all participants to separate WM and RL contributions, we estimated a linear mixed-effects regression with neural RL markers as dependent variable and predictors including behavioral and neural WM components (set size, neural set size markers), RL components (reward history), and their interactions. Shown are means and 95% confidence intervals of fixed effects as well as estimated random effects.

**Table S5. Regression output of model with EEG-SetSize as dependent variable.**

| **Dependent variable: EEG-SetSize** | | | |
|---|---|---|---|
| *Predictors* | *Estimates* | *CI* | *p* |
| (Intercept) | 0 | -0.01 – 0.01 | 0.936 |
| Set Size | 0.12 | 0.10 – 0.13 | **<0.001** |
| Pcor | 0.02 | 0.01 – 0.03 | **<0.001** |
| Set Size × Pcor | 0.01 | 0.00 – 0.02 | **0.015** |
| **Random Effects** | | | |
| $\sigma^2$ | 0.97 | | |
| $\tau_{00\ \text{sbj}}$ | 0 | | |
| $\tau_{11\ \text{sbj.SetSize}}$ | 0 | | |
| $\tau_{11\ \text{sbj.Pcor}}$ | 0 | | |
| $\tau_{11\ \text{sbj.SetSize\_by\_Pcor}}$ | 0 | | |
| $\rho_{01}$ | 0 | | |
| | 0 | | |
| | 0 | | |
| $N_{\text{sbj}}$ | 222 | | |
| Observations | 51794 | | |
| Marginal $R^2$ | 0.014 | | |

Output from multi-linear mixed regression model fitted across all participants. Model included neural set size marker as dependent variable and predictors: set size, reward history (pcor), and their interactions. Shown are means and 95% confidence intervals of fixed effects as well as estimated random effects.

**Table S6. Regression output of model with Accuracy as dependent variable.**

| **Dependent variable: Accuracy** | | | |
|---|---|---|---|
| *Predictors* | *Odds Ratios* | *CI* | *p* |
| (Intercept) | 12.22 | 10.65 – 14.01 | **<0.001** |
| Set size | 0.79 | 0.76 – 0.81 | **<0.001** |
| Neural set size marker | 1.09 | 1.06 – 1.12 | **<0.001** |
| Pcor | 1.71 | 1.67 – 1.75 | **<0.001** |
| Set size × neural set size marker | 1.04 | 1.01 – 1.08 | **0.009** |
| Set size × Pcor | 1.07 | 1.04 – 1.10 | **<0.001** |
| Neural set size marker × Pcor | 1.03 | 1.00 – 1.05 | **0.035** |
| Set size × neural set size marker × Pcor | 1.04 | 1.01 – 1.07 | **0.003** |
| **Random Effects** | | | |
| $\sigma^2$ | 3.29 | | |
| $\tau_{00\ sbj}$ | 1.01 | | |
| $N_{sbj}$ | 222 | | |
| Observations | 58381 | | |
| Marginal $R^2$ | 0.081 | | |

Output from multi-linear mixed regression model fitted across all participants. Model included accuracy as dependent variable and predictors: set size, neural set size marker, reward history (pcor), and their interactions. Shown are means and 95% confidence intervals of fixed effects as well as estimated random effects.

**Table S7. Regression output of model with EEG-RPE as dependent variable.**

| **Dependent variable: EEG-RPE** | | | |
|---|---|---|---|
| *Predictors* | *Estimates* | *CI* | *p* |
| (Intercept) | 0 | -0.01 – 0.01 | 1 |
| Neural Q marker | -0.03 | -0.04 – -0.02 | **<0.001** |
| RPE | -0.07 | -0.09 – -0.06 | **<0.001** |
| **Random Effects** | | | |
| $\sigma^2$ | 0.98 | | |
| $\tau_{00\ \mathrm{sbj}}$ | 0 | | |
| $\tau_{11\ \mathrm{sbj.NeuralQmarker}}$ | 0 | | |
| $\tau_{11\ \mathrm{sbj.RPE}}$ | 0.01 | | |
| $\rho_{01}$ | 0 | | |
| | 0 | | |
| N $_{\mathrm{sbj}}$ | 222 | | |
| Observations | 58381 | | |
| Marginal $R^2$ | 0.006 | | |

Output from multi-linear mixed regression model fitted across all participants. Model included neural reward prediction error (RPE) marker as dependent variable and predictors: RPE (estimated from the RLWM computational model as described in the main manuscript) and neural Q marker. Shown are means and 95% confidence intervals of fixed effects as well as estimated random effects.

**Table S8. Regression output of model with EEG-RPE as dependent variable.**

| | Dependent variable | | |
|---|---|---|---|
| *Predictors* | *Estimates* | *CI* | *p* |
| (Intercept) | 0 | -0.01 – 0.01 | 0.888 |
| Set size | 0.07 | 0.06 – 0.07 | **<0.001** |
| Neural set size marker | -0.01 | -0.02 – -0.00 | **0.036** |
| Set size × neural set size marker | -0.01 | -0.01 – 0.00 | 0.233 |
| **Random Effects** | | | |
| $\sigma^2$ | 0.99 | | |
| $\tau_{00\ sbj}$ | 0 | | |
| $N_{sbj}$ | 222 | | |
| Observations | 51794 | | |
| Marginal $R^2$ | 0.004 | | |

Output from multi-linear mixed regression model fitted across all participants. Model included neural reward prediction error (RPE) marker as dependent variable and predictors: set size, neural set size marker, and their interactions. Shown are means and 95% confidence intervals of fixed effects as well as estimated random effects.

**Table S9. Regression output of clinical model with accuracy as dependent variable.**

| **Dependent variable: Accuracy** | | | |
|---|---|---|---|
| *Predictors* | *Odds Ratios* | *CI* | *p* |
| (Intercept) | 16.24 | 12.90 – 20.45 | **<0.001** |
| Set size | 0.92 | 0.87 – 0.98 | **0.01** |
| Neural set size marker | 1.08 | 1.02 – 1.15 | **0.008** |
| Delay | 0.75 | 0.70 – 0.80 | **<0.001** |
| Pcor | 1.59 | 1.52 – 1.67 | **<0.001** |
| Group [SCZ] | 0.47 | 0.34 – 0.66 | **<0.001** |
| Group [BP] | 0.73 | 0.50 – 1.06 | 0.097 |
| Group [MDD] | 1.17 | 0.81 – 1.68 | 0.402 |
| Set size × neural set size marker | 1.08 | 1.02 – 1.15 | **0.014** |
| Neural set size marker × Delay | 0.94 | 0.88 – 1.00 | **0.043** |
| Neural set size marker × Pcor | 1.01 | 0.96 – 1.06 | 0.665 |
| Delay × Pcor | 1.2 | 1.14 – 1.28 | **<0.001** |
| Set size × Group [SCZ] | 0.92 | 0.85 – 1.00 | **0.05** |
| Set size × Group [BP] | 0.86 | 0.77 – 0.95 | **0.002** |
| Set size × Group [MDD] | 0.95 | 0.85 – 1.05 | 0.285 |
| Neural set size marker × Group [SCZ] | 0.99 | 0.91 – 1.07 | 0.766 |
| Neural set size marker × Group [BP] | 1 | 0.91 – 1.10 | 0.972 |
| Neural set size marker × Group [MDD] | 0.96 | 0.86 – 1.06 | 0.388 |
| Delay × Group [SCZ] | 0.9 | 0.82 – 0.99 | **0.029** |
| Delay × Group [BP] | 0.85 | 0.76 – 0.95 | **0.004** |
| Delay × Group [MDD] | 1.14 | 1.02 – 1.27 | **0.018** |
| Pcor × Group [SCZ] | 0.97 | 0.91 – 1.04 | 0.462 |
| Pcor × Group [BP] | 1 | 0.92 – 1.08 | 0.951 |
| Pcor × Group [MDD] | 1.16 | 1.07 – 1.25 | **<0.001** |
| (Set size × Neural set size marker) × Delay | 1.01 | 0.94 – 1.07 | 0.857 |
| (Set size × Neural set size marker) × Pcor | 1.01 | 0.96 – 1.06 | 0.712 |
| (Set size × Neural set size marker) × Group [SCZ] | 1.01 | 0.93 – 1.10 | 0.832 |
| (Set size × Neural set size marker) × Group [BP] | 0.97 | 0.88 – 1.08 | 0.62 |
| (Set size × Neural set size marker) × Group [MDD] | 1.1 | 0.99 – 1.24 | 0.084 |
| (Neural set size marker × Delay) × Group [SCZ] | 0.96 | 0.88 – 1.05 | 0.422 |
| (Neural set size marker × Delay) × Group [BP] | 0.89 | 0.80 – 0.99 | **0.039** |
| (Neural set size marker × Delay) × Group [MDD] | 1.09 | 0.98 – 1.21 | 0.114 |
| (Neural set size marker × Pcor) × Group [SCZ] | 1.02 | 0.95 – 1.09 | 0.59 |
| (Neural set size marker × Pcor) × Group [BP] | 1.06 | 0.98 – 1.14 | 0.147 |
| (Neural set size marker × Pcor) × Group [MDD] | 0.95 | 0.88 – 1.03 | 0.234 |
| (Delay × Pcor) × Group [SCZ] | 1.06 | 0.98 – 1.15 | 0.141 |
| (Delay × Pcor) × Group [BP] | 1.03 | 0.94 – 1.13 | 0.536 |
| (Delay × Pcor) × Group [MDD] | 1.02 | 0.94 – 1.12 | 0.617 |

*Table S9 continues.*

*Table S9 continued.*

| | | | |
|---|---|---|---|
| (Set size × Neural set size marker × Delay) × Group [SCZ] | 1.1 | 1.01 – 1.20 | **0.0`31** |
| (Set size × Neural set size marker × Delay) × Group [BP] | 1.23 | 1.12 – 1.35 | **<0.001** |
| (Set size × Neural set size marker × Delay) × Group [MDD] | 0.98 | 0.89 – 1.09 | 0.727 |
| (Set size × Neural set size marker × Pcor) × Group [SCZ] | 1.07 | 1.00 – 1.15 | 0.052 |
| (Set size × Neural set size marker × Pcor) × Group [BP] | 0.99 | 0.92 – 1.08 | 0.879 |
| (Set size × Neural set size marker × Pcor) × Group [MDD] | 1.13 | 1.04 – 1.22 | **0.004** |
| **Random Effects** | | | |
| $\sigma^2$ | 3.29 | | |
| $\tau_{00\ sbj:Group}$ | 0.89 | | |
| $\tau_{00}$ Group | 0 | | |
| $N_{sbj}$ | 222 | | |
| N Group | 4 | | |
| Observations | 58381 | | |
| Marginal $R^2$ | 0.169 | | |

Output from multi-linear mixed regression model fitted across all participants and including clinical group (SCZ = schizophrenia, BP = bipolar disorder, MDD = major depressive disorder) as an additional factor. The participants without a mental health disorder diagnosis (CTRL) served as reference group. Regression model included accuracy as binary dependent variable, corrects (1) and errors (0). Predictors included: clinical group, delay, set size, neural set size marker, and previous correct responses indexing reward history (pcor) as well as their interactions. Shown are means and 95% confidence intervals of fixed effects as well as estimated random effects.

**Table S10. Regression output of clinical model with accuracy as dependent variable.**

| **Dependent variable: Accuracy** | | | |
|---|---|---|---|
| *Predictors* | *Odds Ratios* | *CI* | *p* |
| (Intercept) | 19.36 | 16.04 – 23.37 | **<0.001** |
| Set size | 0.9 | 0.84 – 0.96 | **0.001** |
| Delay | 0.71 | 0.66 – 0.76 | **<0.001** |
| Pcor | 1.6 | 1.53 – 1.68 | **<0.001** |
| Group [SCZ] | 0.44 | 0.33 – 0.58 | **<0.001** |
| Group [BP] | 0.68 | 0.50 – 0.93 | **0.016** |
| Group [MDD] | 1.01 | 0.75 – 1.37 | 0.93 |
| Set size × Delay | 0.69 | 0.66 – 0.73 | **<0.001** |
| Set size × Pcor | 0.94 | 0.89 – 0.98 | **0.008** |
| Delay × Pcor | 1.24 | 1.17 – 1.32 | **<0.001** |
| Set size × Group [SCZ] | 0.93 | 0.85 – 1.01 | 0.077 |
| Set size × Group [BP] | 0.87 | 0.79 – 0.97 | **0.01** |
| Set size × Group [MDD] | 0.85 | 0.76 – 0.95 | **0.003** |
| Delay × Group [SCZ] | 0.93 | 0.85 – 1.03 | 0.146 |
| Delay × Group [BP] | 0.91 | 0.81 – 1.02 | 0.108 |
| Delay × Group [MDD] | 1.11 | 0.99 – 1.24 | 0.08 |
| Pcor × Group [SCZ] | 0.99 | 0.93 – 1.05 | 0.742 |
| Pcor × Group [BP] | 0.99 | 0.92 – 1.06 | 0.697 |
| Pcor × Group [MDD] | 1.11 | 1.03 – 1.19 | **0.005** |
| (Set size × Delay) × Group [SCZ] | 1.04 | 0.97 – 1.11 | 0.262 |
| (Set size × Delay) × Group [BP] | 1.02 | 0.94 – 1.11 | 0.633 |
| (Set size × Delay) × Group [MDD] | 1.01 | 0.93 – 1.10 | 0.76 |
| (Set size × Pcor) × Group [SCZ] | 1.08 | 1.00 – 1.15 | **0.039** |
| (Set size × Pcor) × Group [BP] | 1.04 | 0.95 – 1.13 | 0.379 |
| (Set size × Pcor) × Group [MDD] | 1 | 0.92 – 1.08 | 0.951 |
| (Delay × Pcor) × Group [SCZ] | 0.99 | 0.91 – 1.07 | 0.761 |
| (Delay × Pcor) × Group [BP] | 1 | 0.90 – 1.10 | 0.958 |
| (Delay × Pcor) × Group [MDD] | 1 | 0.91 – 1.09 | 0.961 |
| **Random Effects** | | | |
| $\sigma^2$ | 3.29 | | |
| $\tau_{00}$ sbj:Group | 0.69 | | |
| $\tau_{00}$ Group | 0 | | |
| $N_{sbj}$ | 255 | | |
| N Group | 4 | | |
| Observations | 68475 | | |
| Marginal $R^2$ | 0.179 | | |

Output from multi-linear mixed regression model fitted across all participants and including clinical group (SCZ = schizophrenia, BP = bipolar disorder, MDD = major depressive disorder) as an additional factor. The participants without a mental health disorder diagnosis (CTRL) served as reference group. Regression model included accuracy as binary dependent variable, corrects (1) and errors (0). Predictors included: clinical group, delay, set size, and previous correct responses indexing reward history (pcor) as well as their interactions. Shown are means and 95% confidence intervals of fixed effects as well as estimated random effects.

**Table S11. Regression output of clinical model with EEG-SetSize as dependent variable.**

| **Dependent variable: EEG-SetSize** | | | |
|---|---|---|---|
| *Predictors* | *Estimates* | *CI* | *p* |
| (Intercept) | 0.03 | 0.01 – 0.04 | **<0.001** |
| Set size | 0.12 | 0.11 – 0.14 | **<0.001** |
| Group [SCZ] | 0 | -0.03 – 0.02 | 0.711 |
| Group [BP] | 0 | -0.02 – 0.02 | 0.913 |
| Group [MDD] | -0.01 | -0.03 – 0.01 | 0.408 |
| Set size × Pcor | 0.01 | -0.00 – 0.03 | 0.093 |
| Set size × Group [SCZ] | -0.04 | -0.06 – -0.02 | **0.001** |
| Set size × Group [BP] | -0.03 | -0.05 – -0.01 | **0.016** |
| Set size × Group [MDD] | -0.01 | -0.03 – 0.02 | 0.62 |
| Set size × Pcor × Group [SCZ] | 0.01 | -0.02 – 0.03 | 0.622 |
| Set size × Pcor × Group [BP] | 0 | -0.03 – 0.02 | 0.822 |
| Set size × Pcor × Group [MDD] | -0.01 | -0.04 – 0.01 | 0.232 |
| **Random Effects** | | | |
| $\sigma^2$ | 0.98 | | |
| $\tau_{00\ sbj}$ | 0 | | |
| $N_{sbj}$ | 222 | | |
| Observations | 58381 | | |
| Marginal $R^2$ | 0.012 | | |

Output from multi-linear mixed regression model fitted across all participants and including clinical group (SCZ = schizophrenia, BP = bipolar disorder, MDD = major depressive disorder) as an additional factor. The participants without a mental health disorder diagnosis (CTRL) served as reference group. Regression model included neural set size marker as dependent variable. Predictors included: clinical group, set size, and its interaction with previous correct responses indexing reward history (pcor). Shown are means and 95% confidence intervals of fixed effects as well as estimated random effects.

**Table S12. Regression output of clinical model with EEG-Q as dependent variable.**

| **Dependent variable: EEG-Q** | | | |
|---|---|---|---|
| *Predictors* | *Estimates* | *CI* | *p* |
| (Intercept) | 0 | -0.02 – 0.02 | 0.816 |
| Pcor | 0.13 | 0.11 – 0.14 | **<0.001** |
| Neural set size marker | -0.31 | -0.33 – -0.30 | **<0.001** |
| Set size | 0.07 | 0.05 – 0.09 | **<0.001** |
| Group [SCZ] | 0 | -0.03 – 0.03 | 0.964 |
| Group [BP] | 0 | -0.03 – 0.03 | 0.889 |
| Group [MDD] | 0.01 | -0.02 – 0.04 | 0.667 |
| Pcor × Neural set size marker | 0.02 | 0.00 – 0.03 | **0.02** |
| Pcor × Set size | 0 | -0.01 – 0.02 | 0.581 |
| Neural set size marker × Set size | 0 | -0.01 – 0.02 | 0.861 |
| Pcor × Group [SCZ] | -0.02 | -0.05 – 0.00 | 0.052 |
| Pcor × Group [BP] | -0.01 | -0.04 – 0.01 | 0.395 |
| Pcor × Group [MDD] | 0 | -0.02 – 0.02 | 0.937 |
| Neural set size marker × Group [SCZ] | -0.03 | -0.05 – -0.01 | **0.005** |
| Neural set size marker × Group [BP] | -0.03 | -0.06 – -0.01 | **0.004** |
| Neural set size marker × Group [MDD] | -0.02 | -0.04 – 0.00 | 0.086 |
| Set size × Group [SCZ] | -0.01 | -0.04 – 0.03 | 0.673 |
| Set size × Group [BP] | -0.05 | -0.08 – -0.01 | **0.014** |
| Set size × Group [MDD] | -0.01 | -0.05 – 0.02 | 0.485 |
| (Pcor × Neural set size marker) × Set size | -0.01 | -0.02 – 0.01 | 0.212 |
| (Pcor × Neural set size marker) × Set size × Group [SCZ] | -0.02 | -0.04 – 0.00 | 0.104 |
| (Pcor × Neural set size marker) × Set size × Group [BP] | 0 | -0.02 – 0.02 | 0.931 |
| (Pcor × Neural set size marker) × Set size × Group [MDD] | -0.01 | -0.03 – 0.01 | 0.424 |
| (Pcor × Set size) × Group [SCZ] | -0.01 | -0.03 – 0.02 | 0.489 |
| (Pcor × Set size) × Group [BP] | 0 | -0.03 – 0.02 | 0.835 |
| (Pcor × Set size) × Group [MDD] | 0.01 | -0.01 – 0.04 | 0.299 |
| (Neural set size marker × Set size) × group [SCZ] | 0 | -0.03 – 0.02 | 0.732 |
| (Neural set size marker × Set size) × group [BP] | 0 | -0.03 – 0.02 | 0.721 |
| (Neural set size marker × Set size) × group [MDD] | -0.01 | -0.04 – 0.01 | 0.255 |
| (Pcor × neural set size marker × Set size) × Group [SCZ] | 0.01 | -0.01 – 0.03 | 0.366 |
| (Pcor × neural set size marker × Set size) × Group [BP] | 0 | -0.03 – 0.02 | 0.846 |
| (Pcor × neural set size marker × Set size) × Group [MDD] | 0.01 | -0.01 – 0.03 | 0.36 |
| **Random Effects** | | | |
| $\sigma^2$ | 0.86 | | |
| $\tau_{00\ SetSize}$ | 0.01 | | |
| $\tau_{00\ sbj}$ | 0 | | |
| $\tau_{11\ SetSize.Pcor}$ | 0 | | |
| $\tau_{11\ sbj.SetSize}$ | 0 | | |

*Table S12 continues.*

*Table S12 continued.*

| | |
|---|---|
| $\rho_{01\ SetSize}$ | 0.22 |
| $\rho_{01\ sbj}$ | 0 |
| $N_{\ sbj}$ | 222 |
| $N_{\ SetSize}$ | 888 |
| Observations | 51794 |
| Marginal $R^2$ | 0.124 |

Output from multi-linear mixed regression model fitted across all participants and including clinical group (SCZ = schizophrenia, BP = bipolar disorder, MDD = major depressive disorder) as an additional factor. The participants without a mental health disorder diagnosis (CTRL) served as reference group. Regression model included neural Q marker as dependent variable. Predictors included: clinical group, set size, neural set size marker, previous correct responses indexing reward history (pcor), and their interactions. Shown are means and 95% confidence intervals of fixed effects as well as estimated random effects.

**Table S13. Regression output of clinical model with EEG-SetSize as dependent variable.**

| **Dependent variable: EEG-SetSize** | | | |
|---|---|---|---|
| *Predictors* | *Estimates* | *CI* | *p* |
| (Intercept) | 0.03 | 0.01 – 0.04 | **<0.001** |
| Set size | 0.15 | 0.13 – 0.16 | **<0.001** |
| Delay | -0.03 | -0.05 – -0.02 | **<0.001** |
| Pcor | 0.03 | 0.01 – 0.04 | **<0.001** |
| Group [SCZ] | -0.01 | -0.03 – 0.02 | 0.673 |
| Group [BP] | 0 | -0.03 – 0.02 | 0.756 |
| Group [MDD] | -0.01 | -0.04 – 0.01 | 0.268 |
| Set size × Delay | 0.02 | 0.00 – 0.03 | **0.049** |
| Set size × Pcor | 0 | -0.01 – 0.02 | 0.583 |
| Delay × Pcor | 0.01 | -0.01 – 0.02 | 0.251 |
| Set size × Group [SCZ] | -0.03 | -0.05 – -0.00 | **0.021** |
| Set size × Group [BP] | -0.02 | -0.04 – 0.01 | 0.192 |
| Set size × Group [MDD] | 0 | -0.03 – 0.02 | 0.722 |
| Delay × Group [SCZ] | -0.02 | -0.04 – 0.01 | 0.202 |
| Delay × Group [BP] | -0.03 | -0.05 – -0.00 | **0.039** |
| Delay × Group [MDD] | 0 | -0.03 – 0.02 | 0.693 |
| Pcor × Group [SCZ] | 0 | -0.03 – 0.02 | 0.681 |
| Pcor × Group [BP] | 0 | -0.02 – 0.03 | 0.838 |
| Pcor × Group [MDD] | -0.03 | -0.05 – -0.00 | **0.032** |
| (Set size × Delay) × Group [SCZ] | 0.01 | -0.01 – 0.04 | 0.302 |
| (Set size × Delay) × Group [BP] | 0.01 | -0.01 – 0.04 | 0.295 |
| (Set size × Delay) × Group [MDD] | 0 | -0.02 – 0.03 | 0.698 |
| (Set size × Pcor) × Group [SCZ] | 0.02 | -0.01 – 0.04 | 0.149 |
| (Set size × Pcor) × Group [BP] | 0 | -0.02 – 0.03 | 0.819 |
| (Set size × Pcor) × Group [MDD] | 0 | -0.02 – 0.02 | 0.955 |
| (Delay × Pcor) × Group [SCZ] | 0 | -0.03 – 0.02 | 0.822 |
| (Delay × Pcor) × Group [BP] | 0 | -0.03 – 0.02 | 0.698 |
| (Delay × Pcor) × Group [MDD] | 0 | -0.03 – 0.02 | 0.761 |
| **Random Effects** | | | |
| $\sigma^2$ | 0.97 | | |
| $\tau_{00\ sbj}$ | 0 | | |
| $N_{sbj}$ | 222 | | |
| Observations | 51794 | | |
| Marginal $R^2$ | 0.017 | | |

Output from multi-linear mixed regression model fitted across all participants and including clinical group (SCZ = schizophrenia, BP = bipolar disorder, MDD = major depressive disorder) as an additional factor. The participants without a mental health disorder diagnosis (CTRL) served as reference group. Regression model included neural set size marker as dependent variable. Predictors included: clinical group, set size, delay, previous correct responses indexing reward history (pcor), and their two-way interactions with group. Shown are means and 95% confidence intervals of fixed effects as well as estimated random effects.

**Table S14. Regression output of clinical model with EEG-SetSize as dependent variable.**

| **Dependent variable: EEG-SetSize** | | | |
|---|---|---|---|
| *Predictors* | *Estimates* | *CI* | *p* |
| (Intercept) | 0.04 | 0.02 – 0.05 | **<0.001** |
| Set size | 0.13 | 0.11 – 0.15 | **<0.001** |
| Group [SCZ] | 0 | -0.02 – 0.02 | 0.989 |
| Group [BP] | 0 | -0.02 – 0.03 | 0.849 |
| Group [MDD] | -0.01 | -0.04 – 0.01 | 0.305 |
| Set size × Group [SCZ] | -0.04 | -0.07 – -0.01 | **0.014** |
| Set size × Group [BP] | -0.04 | -0.07 – -0.00 | **0.037** |
| Set size × Group [MDD] | -0.01 | -0.04 – 0.02 | 0.661 |
| **Random Effects** | | | |
| $\sigma^2$ | 0.97 | | |
| $\tau_{00\ sbj}$ | 0 | | |
| $\tau_{11\ sbj.SetSize}$ | 0 | | |
| $\rho_{01\ sbj}$ | -1 | | |
| N $_{sbj}$ | 222 | | |
| Observations | 51794 | | |
| Marginal $R^2$ | 0.014 | | |

Output from multi-linear mixed regression model fitted across all participants and including clinical group (SCZ = schizophrenia, BP = bipolar disorder, MDD = major depressive disorder) as an additional factor. The participants without a mental health disorder diagnosis (CTRL) served as reference group. Regression model included neural set size marker as dependent variable. Predictors included: clinical group, set size and their interactions. Shown are means and 95% confidence intervals of fixed effects as well as estimated random effects.

**Table S15. Regression output of clinical model with EEG-RPE as dependent variable.**

| **Dependent variable: EEG-RPE** | | | |
|---|---|---|---|
| *Predictors* | *Estimates* | *CI* | *p* |
| (Intercept) | -0.16 | -0.19 – -0.14 | **<0.001** |
| Set size | 0.06 | 0.04 – 0.08 | **<0.001** |
| Neural set size marker | 0.02 | -0.00 – 0.03 | 0.105 |
| Group [SCZ] | 0.05 | 0.02 – 0.09 | **0.002** |
| Group [BP] | 0.05 | 0.01 – 0.09 | **0.01** |
| Group [MDD] | -0.01 | -0.05 – 0.03 | 0.629 |
| Set size × neural set size marker | 0 | -0.02 – 0.01 | 0.831 |
| Set size × Group [SCZ] | 0.02 | -0.01 – 0.05 | 0.267 |
| Set size × Group [BP] | 0.02 | -0.02 – 0.06 | 0.252 |
| Set size × Group [MDD] | 0 | -0.04 – 0.03 | 0.937 |
| Neural set size marker × Group [SCZ] | -0.05 | -0.07 – -0.02 | **0.001** |
| Neural set size marker × Group [BP] | -0.04 | -0.08 – -0.01 | **0.005** |
| Neural set size marker × Group [MDD] | -0.02 | -0.05 – 0.00 | **0.049** |
| (Set size × Neural set size marker) × Group [SCZ] | -0.03 | -0.05 – -0.01 | **0.014** |
| (Set size × Neural set size marker) × Group [BP] | 0 | -0.03 – 0.02 | 0.911 |
| (Set size × Neural set size marker) × Group [MDD] | 0.01 | -0.01 – 0.04 | 0.287 |
| **Random Effects** | | | |
| $\sigma^2$ | 0.95 | | |
| $\tau_{00\ sbj}$ | 0.01 | | |
| $\tau_{11\ sbj.SetSize}$ | 0.01 | | |
| $\tau_{11\ sbj.NeuralSetSizeMarker}$ | 0 | | |
| $\tau_{11\ sbj.SetSize:NeuralSetSizeMarker}$ | 0 | | |
| $\rho_{01}$ | 0.04 | | |
| | 0 | | |
| | 0.03 | | |
| $N_{sbj}$ | 222 | | |
| Observations | 51794 | | |
| Marginal $R^2$ | 0.006 | | |

Output from multi-linear mixed regression model fitted across all participants and including clinical group (SCZ = schizophrenia, BP = bipolar disorder, MDD = major depressive disorder) as an additional factor. The participants without a mental health disorder diagnosis (CTRL) served as reference group. Regression model included neural reward prediction error (RPE) as dependent variable. Predictors included: clinical group, set size, neural set size marker, and their interactions. Shown are means and 95% confidence intervals of fixed effects as well as estimated random effects.

**Table S16. Regression output of clinical model with EEG-RPE as dependent variable.**

| **Dependent variable: EEG-RPE** | | | |
|---|---|---|---|
| *Predictors* | *Estimates* | *CI* | *p* |
| (Intercept) | -0.18 | -0.20 – -0.15 | **<0.001** |
| Set size | 0.05 | 0.04 – 0.07 | **<0.001** |
| Delay | 0.03 | 0.02 – 0.05 | **<0.001** |
| Pcor | -0.16 | -0.17 – -0.14 | **<0.001** |
| Group [SCZ] | 0.04 | 0.01 – 0.08 | **0.009** |
| Group [BP] | 0.04 | 0.00 – 0.07 | **0.048** |
| Group [MDD] | -0.01 | -0.05 – 0.02 | 0.5 |
| Set size × Delay | 0.02 | 0.01 – 0.04 | **0.001** |
| Set size × Pcor | -0.02 | -0.03 – -0.00 | **0.018** |
| Delay × Pcor | -0.05 | -0.06 – -0.03 | **<0.001** |
| Set size × Group [SCZ] | 0.02 | -0.00 – 0.04 | 0.102 |
| Set size × Group [BP] | 0.03 | -0.00 – 0.05 | 0.058 |
| Set size × Group [MDD] | 0 | -0.02 – 0.03 | 0.86 |
| Delay × Group [SCZ] | 0 | -0.03 – 0.02 | 0.782 |
| Delay × Group [BP] | 0 | -0.03 – 0.02 | 0.983 |
| Delay × Group [MDD] | 0 | -0.02 – 0.02 | 0.985 |
| Pcor × Group [SCZ] | 0 | -0.02 – 0.02 | 0.939 |
| Pcor × Group [BP] | -0.03 | -0.05 – -0.00 | **0.022** |
| Pcor × Group [MDD] | -0.03 | -0.05 – -0.01 | **0.006** |
| (Set size × Delay) × Group [SCZ] | 0.01 | -0.01 – 0.03 | 0.447 |
| (Set size × Delay) × Group [BP] | 0.03 | 0.00 – 0.05 | **0.036** |
| (Set size × Delay) × Group [MDD] | 0.01 | -0.01 – 0.04 | 0.265 |
| (Set size× Pcor) × Group [SCZ] | -0.01 | -0.03 – 0.02 | 0.537 |
| (Set size× Pcor) × Group [BP] | 0.01 | -0.01 – 0.04 | 0.354 |
| (Set size× Pcor) × Group [MDD] | -0.01 | -0.04 – 0.01 | 0.293 |
| (Delay × Pcor) × Group [SCZ] | 0.01 | -0.01 – 0.03 | 0.362 |
| (Delay × Pcor) × Group [BP] | 0 | -0.02 – 0.03 | 0.769 |
| (Delay × Pcor) × Group [MDD] | -0.02 | -0.04 – 0.00 | 0.053 |
| **Random Effects** | | | |
| $\sigma^2$ | 0.93 | | |
| $\tau_{00\ sbj}$ | 0 | | |
| N $_{sbj}$ | 222 | | |
| Observations | 51794 | | |
| Marginal $R^2$ | 0.041 | | |

Output from multi-linear mixed regression model fitted across all participants and including clinical group (SCZ = schizophrenia, BP = bipolar disorder, MDD = major depressive disorder) as an additional factor. The participants without a mental health disorder diagnosis (CTRL) served as reference group. Regression model included neural reward prediction error (RPE) as dependent variable. Predictors included: clinical group, set size, delay, previous correct responses indexing reward history (pcor), and their two-way interactions with group. Shown are means and 95% confidence intervals of fixed effects as well as estimated random effects.
